# Early prediction of preeclampsia using the first trimester vaginal microbiome

**DOI:** 10.1101/2024.12.01.626267

**Authors:** William F. Kindschuh, George I. Austin, Yoli Meydan, Heekuk Park, Julia A. Urban, Emily Watters, Susan Pollak, George R. Saade, Judith Chung, Brian M. Mercer, William A. Grobman, David M. Haas, Robert M. Silver, Myrna Serrano, Gregory A. Buck, Rebecca McNeil, Renu Nandakumar, Uma Reddy, Ronald J. Wapner, Aya Brown Kav, Anne-Catrin Uhlemann, Tal Korem

**Author notes:** These authors contributed equally and are randomly ordered (William F. Kindschuh, George I. Austin).

## Abstract

Preeclampsia is a severe obstetrical syndrome which contributes to 10-15% of all maternal deaths. Although the mechanisms underlying systemic damage in preeclampsia—such as impaired placentation, endothelial dysfunction, and immune dysregulation—are well studied, the initial triggers of the condition remain largely unknown. Furthermore, although the pathogenesis of preeclampsia begins early in pregnancy, there are no early diagnostics for this life-threatening syndrome, which is typically diagnosed much later, after systemic damage has already manifested. Here, we performed deep metagenomic sequencing and multiplex immunoassays of vaginal samples collected during the first trimester from 124 pregnant individuals, including 62 who developed preeclampsia with severe features. We identified multiple significant associations between vaginal immune factors, microbes, clinical factors, and the early pathogenesis of preeclampsia. These associations vary with BMI, and stratification revealed strong associations between preeclampsia and *Bifidobacterium* spp., *Prevotella timonensis*, and *Sneathia vaginalis*. Finally, we developed machine learning models that predict the development of preeclampsia using this first trimester data, collected ~5.7 months prior to clinical diagnosis, with an auROC of 0.78. We validated our models using data from an independent cohort (MOMS-PI), achieving an auROC of 0.80. Our findings highlight robust associations among the vaginal microbiome, local host immunity, and early pathogenic processes of preeclampsia, paving the way for early detection, prevention and intervention for this devastating condition.

## Introduction

Preeclampsia is a multi-system hypertensive disorder that complicates approximately 5% of all pregnancies, leading to 10-15% of maternal deaths and up to 25% of neonatal deaths^1–3^. It also increases the risk for additional adverse outcomes such as intrauterine growth restriction and preterm delivery^4^. The pathogenesis of preeclampsia begins early in pregnancy, as the placental vasculature fails to remodel properly, leading to poor placental perfusion^4^. This is followed later in pregnancy by the release of circulating factors from the placenta, which trigger systemic endothelial dysfunction, and increase the risk for maternal morbidities such as seizures, stroke, and hemorrhage^4^.

The identification of individuals early in pregnancy who are at high risk for developing preeclampsia would enable researchers to more effectively study the early pathogenesis of this syndrome, improve utilization of existing preventative therapies^5^, and facilitate the development of novel treatments^6^. However, although the pathology of preeclampsia begins in the first trimester, it is currently diagnosed only late in pregnancy. This is, by definition, as diagnosis is currently based on the presence of maternal hypertension after 20 weeks of gestation combined with additional symptoms such as proteinuria, vision changes, headaches, and elevated transaminase levels^4,7^, which manifest late in the disease and indicate impending organ failure. This delayed diagnosis severely limits opportunities for early intervention. Several recent efforts have been made to improve the early diagnosis of preeclampsia. For example, a diagnostic test based on the ratio of serum soluble fms-like tyrosine kinase 1 (sFlt-1) and placental growth factor (PlGF) has achieved an area under the receiver operating characteristic curve (auROC) of 0.82 for predicting a preeclampsia diagnosis within 4 weeks in individuals already identified as high-risk for the syndrome^8^. Similar efforts have been made to develop a diagnostic test for preeclampsia based on cell-free RNA levels in maternal serum in the second trimester, also achieving an auROC of 0.82^9^.

Maternal immune dysregulation, which has been repeatedly observed in individuals with preeclampsia^10^, offers both a potential avenue for the development of novel diagnostics and a window into the pathogenesis of preeclampsia early in pregnancy. A systemic increase in serum CD4+ T cells^11^, circulating proinflammatory cytokines^12^, and a decrease in regulatory T cells^13,14^ are all associated with preeclampsia. Other studies have found cytokine levels^15,16^ and immune cell populations^17^ to be altered in the placentas of individuals with preeclampsia, albeit with measurements taken postpartum. Altogether, these studies suggest that both local and systemic immune dysregulation may be implicated in the pathogenesis of preeclampsia.

The involvement of a microbial trigger in the pathogenesis of preeclampsia has been suggested previously^18,19^ and specifically for periodontal and urinary tract infections^20^. Another potential trigger is the vaginal microbiome, which has been highlighted as a source of ascending infection during pregnancy^21^, was shown to promote inflammation in the reproductive tract^22–24^, disrupt the cervicovaginal epithelial barrier^23^, and was associated with other adverse pregnancy outcomes, such as preterm premature rupture of membranes^25,26^. Two recent studies have found associations between preeclampsia and the vaginal microbiome sampled in the third trimester, after preeclampsia was diagnosed^27,28^. However, studies that investigate the vaginal microbiome early in pregnancy are still needed to advance our understanding of the role of this ecosystem during the early pathogenesis of preeclampsia, and to evaluate the potential for early diagnostics.

Here, we investigate the early pregnancy (6+0-13+6 weeks+days of gestation) vaginal microbiome and host immunity and their interaction with preeclampsia with severe features and related clinical factors in 124 individuals. We identified several associations between the composition of the vaginal microbiome, levels of vaginal immune factors, and the development of preeclampsia, and demonstrated that some of these associations are impacted by maternal BMI. We then developed predictive models for preeclampsia using this data, which we validated in an independent pregnancy cohort. The performance and generalizability of our models demonstrate a strong association of the early pregnancy vaginal microbiome and local immunity with the development of preeclampsia.

## Results

### Vaginal microbiota, immune factors, and clinical data from a multi-center pregnancy cohort

To study the interaction of the vaginal microbiome and immune system with the development of preeclampsia we compiled a subcohort of the nuMoM2b cohort^29^, a large multi-center observational study of nulliparous pregnant individuals with singleton gestations (**Fig. 1**; **Methods**). As preeclampsia with severe features (sPEC) accounts for over one third of all cases of preeclampsia^30^ and a majority of preeclampsia related morbidity and mortality^31,32^, we focused our investigation on this more extreme phenotype, and randomly selected 62 individuals who developed sPEC and 62 who were not diagnosed with sPEC, but could have other hypertensive disorders, including mild preeclampsia (**Table 1; Methods**; hereafter termed “non-sPEC”). Both groups were frequency matched on maternal age, race, and clinical enrollment site (**Methods**). The nuMoM2b study collected extensive clinical data and medical history, including first trimester maternal BMI, age, hypertension status^29^. Consistent with prior research^33,34^, individuals who developed sPEC had higher BMI (Mann-Whitney *U p*=0.03), higher first trimester blood pressure (*p*=5.8×10^−5^), and gave birth earlier in pregnancy (*p*=0.0006; **Table 1**).

**Figure 1.**
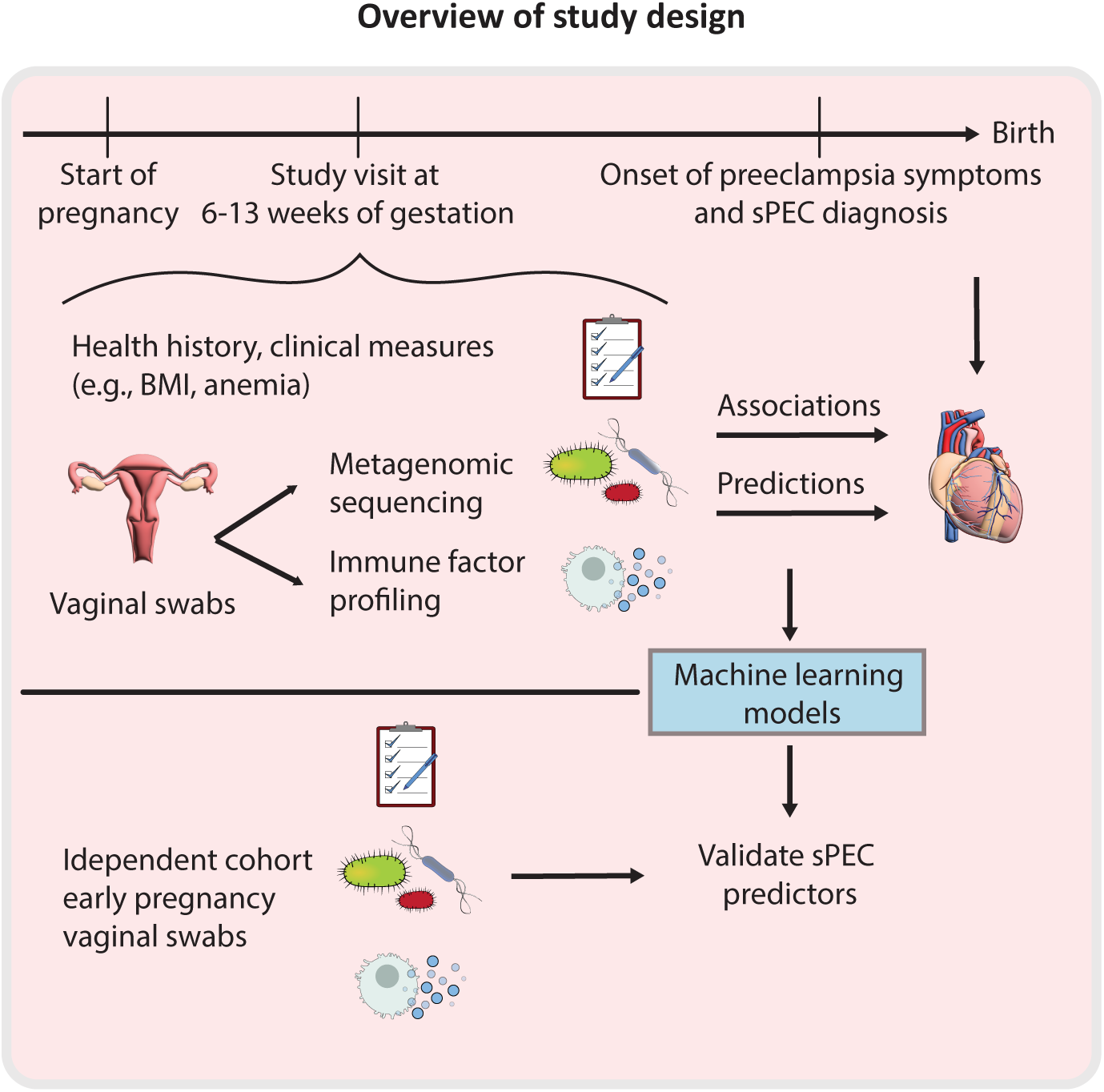
Study overview.

**Table 1.** Cohort characteristics. sPEC, preeclampsia with severe features; BMI, body mass index; GA, gestational age; *p* - Fisher’s exact or Mann-Whitney *U* test, as appropriate.

|  | sPEC | non-sPEC | Difference<br>(*p* value) |
| --- | --- | --- | --- |
| **N** | 62 | 62 |  |
| **Race / ethnicity (N (%))** |  |  | 0.92 |
| **Non-Hispanic Black** | 15 (24.2) | 15 (24.2) | 1.00 |
| **Non-Hispanic White** | 26 (41.9) | 26 (41.9) | 1.00 |
| **Hispanic** | 12 (19.4) | 10 (16.1) | 0.81 |
| **Asian** | 4 (6.5) | 3 (4.8) | 1.00 |
| **Other** | 5 (8.1) | 8 (12.9) | 0.56 |
| **Mean arterial blood pressure at first trimester (mean mmHg ±sd)** | 87±10 | 80±8 | 5.8x10<sup>-5</sup> |
| **GA at delivery (median weeks [range])** | 38.3<br>[24.4–42.1] | 39.5<br>[21.1–41.9] | 0.0006 |
| **BMI at first trimester (median kg/m<sup>2</sup> [IQR])** | 28.2<br>[24.6–34.1] | 25.8 [22.2–30.7] | 0.029 |
| **Age (mean years±sd)** | 26.6±5.9 | 25.9±5.9 | 0.58 |
| **GA at sPEC diagnosis (median weeks [range])** | 38.0<br>[23.9–1 week postpartum] | -- | -- |
| **Other hypertensive disorders (N (%))** |  |  |  |
| **Antepartum hypertension** |  | 3 (4.8) |  |
| **Intra or postpartum hypertension** |  | 4 (6.5) |  |
| **Mild preeclampsia** |  | 2 (3.2) |  |

We analyzed vaginal swabs which were self-collected between 6.4 – 13.9 weeks of gestation (mean±sd 11.9±1.6) of pregnancy, 2.4–7.6 months before sPEC diagnosis (mean±sd 5.7±1.0 months). We performed multiplex immunoassays (Luminex) to measure the levels of 20 pro- and anti-inflammatory immune factors: EGF, Flt-3L, G-CSF, IFNα2, IFN-γ, IL-10, IL-18, IL-1RA, IL-1α, IL-1β, IL-27, IL-4, IL-6, IL-8, MCP-1, RANTES, TGFα, TNFα, VEGF, and sCD40L (**Methods**). We selected these based on the ability to obtain measurements in our samples as well as review of existing literature regarding vaginal immune factors in pregnancy^35–39^. We also profiled the vaginal microbiome via shotgun metagenomic sequencing to a mean±sd depth of 43.23±23.4 million reads, yielding an average of 5.83±7.6 million microbial reads after processing and discarding reads classified as human DNA (**Methods**).

### Early pregnancy vaginal immune factors are associated with sPEC

To determine whether the vaginal immune profile in early pregnancy is associated with the development of sPEC, we compared the overall profile of individuals who developed sPEC to those who did not. We found that overall immune factor levels were weakly associated with the development of sPEC (PERMANOVA *p*=0.029; **Fig. 2a**), driven by their first principal component (PC1), which captured 32% of the variance (Mann-Whitney *U p*=0.013; **Fig. 2b**). No individual immune factor dominated the composition of this first principal component, with all immune factors having coefficients between 0.13 and 0.33, except EGF which had a negative coefficient (**Fig. S1**). These results demonstrate that the association of the immune system with the early pathogenesis of sPEC is attributable to variation in multiple immune factors.

**Figure 2.**
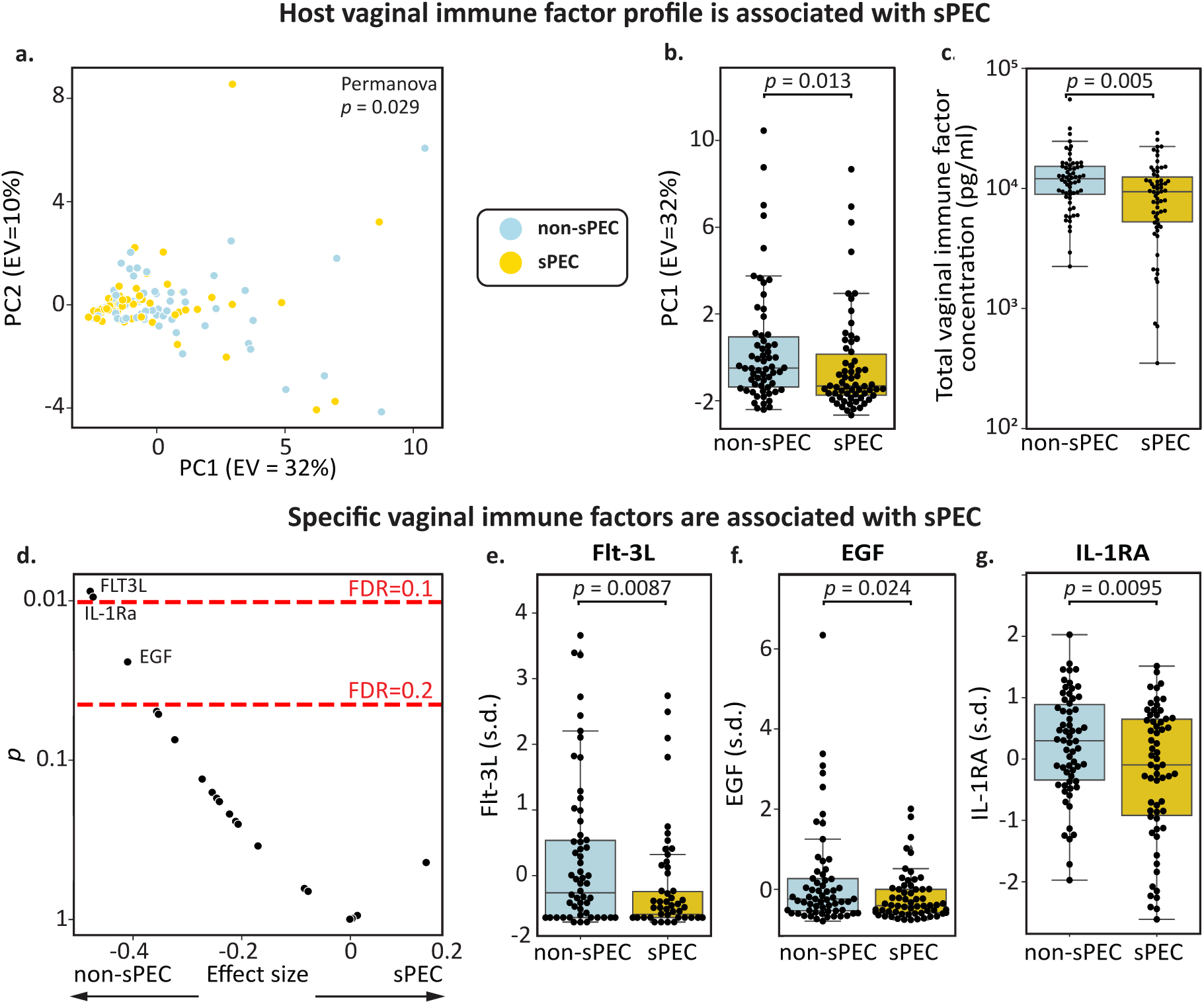
Vaginal immune factors are associated with sPEC. **a,** PCA of immune factor profiles colored by future sPEC status (PERMANOVA *p*=0.029). **b**, Box and swarm plot demonstrating a significant association between immune factor PC1 and sPEC (Mann-Whitney *U p*=0.013). **c**, Box and swarm plot demonstrating a significant association between the total concentration of the 20 immune factors that we measured and sPEC (Mann-Whitney *U p*=0.005). **d**, Significance (y-axis; two-sided t-test) and effect size (x-axis; Cohen’s d) for associations between individual immune factors and sPEC. Most immune factors tested have lower vaginal levels in individuals who developed sPEC, consistent with studies of placental immune factors in PEC^16^ and vaginal immune factors in preterm birth^40^. **e-g**, Box and swarm plots of Flt-3L (e), EGF (f) and IL-1RA (g), all of which were significantly depleted in the vaginal ecosystem of individuals who developed sPEC (*q*=0.09, 0.16, 0.09, respectively). Box, IQR; line, median; whiskers, nearest point to 1.5*IQR.

Intriguingly, we found a significant reduction in the total concentration of immune factors measured in this group (Mann-Whiteny *U p*=0.005, **Fig. 2c**). When next investigating the associations of specific immune factors with sPEC, we found that this pattern was consistent across most immune factors, as nearly all had lower levels in individuals who developed sPEC (17 out of 20; **Fig. 2d**). Similar reductions in both pro- and anti-inflammatory immune factors in PEC were previously reported in the placenta^16^, and a similar observation of lower levels was also reported in the vaginal ecosystem for other adverse outcomes, such as spontaneous preterm birth^40^. Therefore, our results suggest a phenomenon of reduced local immune factor concentrations in sPEC and potentially other pathologies.

We identified three specific immune factors which were significantly reduced in individuals who subsequently developed sPEC (t-test *p*<0.05, *q*<0.2; **Fig. 2d**): Flt-3L (*p*=0.0087, *q*=0.095; **Fig. 2e**), a pro-inflammatory cytokine that activates dendritic cells in response to infection^41,42^; EGF (*p*=0.024, *q*=0.16, **Fig. 2f**), a growth factor that stimulates the proliferation of epithelial cells and fibroblasts^43,44^; and IL-1RA (*p*=0.0095, *q*=0.095; **Fig. 2g**), a competitive inhibitor of the IL-1 receptor with a powerful anti-inflammatory effect^45,46^. While it has similarly been reported that placental EGF is depleted in individuals with PEC^47^, in the serum, levels of IL-1RA and Flt-3L late in pregnancy have previously been shown to be positively associated with PEC^48,49^, suggesting that associations between immune factors and sPEC may be influenced by gestational age or biological niche. Overall, our results suggest that the local immune system in the vagina may be dysregulated early in pregnancy, months before symptoms become evident, among individuals who develop sPEC.

### The early pregnancy vaginal microbiome is associated with the development of sPEC

We next investigated the association of vaginal microbes in early pregnancy with the development of sPEC. To this end, we quantified strain level abundances (ANI=99%) using a reference derived from a recent large-scale assembly effort^50^ (**Methods**). Similarly to our analysis of immune factors, we first used robust principal component analysis^51^ to examine the difference between the entire microbiome profiles of our cohort (**Methods**). However, we did not detect a significant association between the vaginal microbiome and sPEC (PERMANOVA *p*=0.12; **Fig. 3a**) or between the first microbiome robust principal component (RPC1) and sPEC (Mann-Whitney *U p*=0.21; **Fig. 3b**). As BMI is associated with both PEC risk^4,33^ and the vaginal microbiome^52^, we examined microbiome-sPEC associations stratified by first trimester BMI. We identified a relatively stronger association between the vaginal microbiome and sPEC in individuals with higher BMI (BMI≥25; N=67, N sPEC=37), both when examining the entire microbiome profile (PERMANOVA *p*=0.01; **Fig. 3a**) and when examining only RPC1 (Mann-Whitney *U p*=0.021; **Fig. 3b**). While we did not detect statistically significant associations in individuals with lower BMI (BMI<25; N=43, N sPEC=20; PERMANOVA *p* = 0.46; **Fig. 3a**; *p*=0.39; **Fig. 3b**), we noted opposite trends in the first principal coordinate between the two groups (higher values for sPEC in BMI<25, while lower in BMI≥25; **Fig. 3b**). When examining whether community state types (CSTs), measured as dominance with >30% abundance^22^ (or lack thereof; **Methods**), we found a weak association between *Gardnerella* dominance and non-sPEC among individuals with BMI≥25 (Fisher’s exact test *p*=0.03, *q*=0.14), but not in other groups or CSTs (**Fig. S2a-b**). These different trends suggest that BMI, which is associated with metabolic health and systemic inflammation^53^, may impact the associations between vaginal microbes and sPEC. This highlights the need for considering the vaginal microbiome within a broader clinical context when investigating its associations with sPEC.

**Figure 3.**
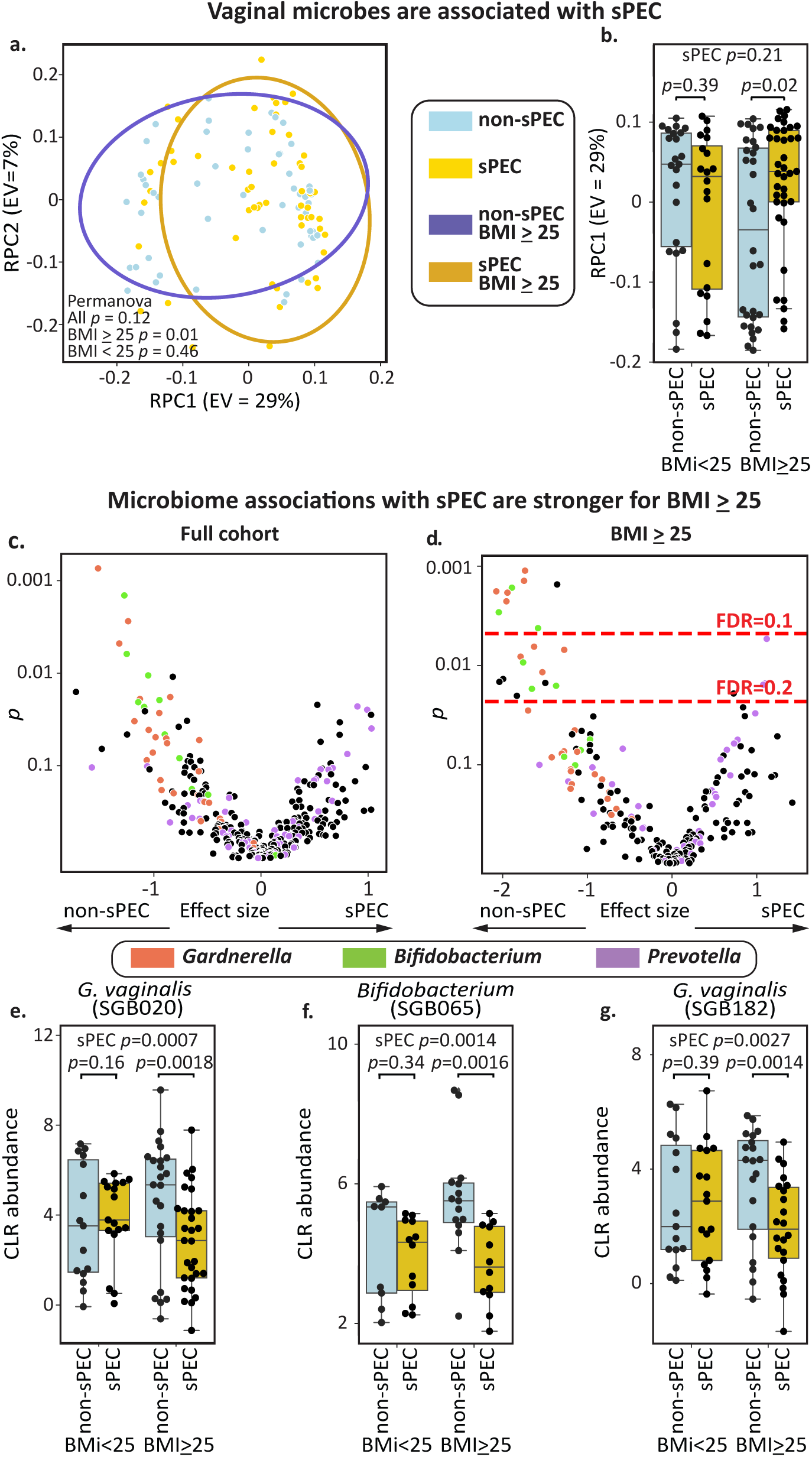
Abundance of early pregnancy vaginal microbes are associated with sPEC development. **a,** Scatterplot of the first two robust principal components^51^ of the vaginal microbiome, colored by subsequent sPEC diagnosis; ellipses, covariance error. **b**, Box and swarm plots of the first principal component of the vaginal microbiome, stratified by BMI group and sPEC. An association between the vaginal microbiome and sPEC is observed only in individuals with higher BMI, as there are opposite trends between the BMI groups. **c,** Significance (y-axis; two-sided Mann-Whitney *U* test of CLR-transformed values) and effect size (x-axis; difference in mean) for associations between specific vaginal strains (**Methods**). **d,** Similar to c, but evaluated within individuals with BMI≥25. Dashed red lines indicate FDR thresholds of 0.2 and 0.1. **e-g,** Box and swarm plots of the three vaginal taxa with the most significant associations with sPEC within individuals with BMI≥25, stratified by BMI group and sPEC. Box, IQR; line, median; whiskers, nearest point to 1.5*IQR; *p*, Mann-Whitney *U* test.

We next investigated associations between specific microbes and the development of sPEC. Once again, we only identified weak associations in the full cohort (no taxa with Mann-Whitney *U q*<0.2; **Fig. 3c**). Additionally, while we did not find any differentially abundant or dominant taxa in individuals with BMI<25 (Mann-Whitney *U p*>0.01, *q>*0.2 for all taxa; **Fig. S2c-d**), likely due in part to smaller sample size, we identified 24 taxa that were significantly associated with the development of sPEC in the high BMI group (Mann-Whitney *U p*<0.05, *q*<0.1 for 9 taxa, 0.1<*q*<0.2 for 15; **Fig. 3d**). The top three taxa with the most significant associations with sPEC were two *G. vaginalis* strains (*p* = 0.0007 overall, *p* = 0.0018 within BMI≥25 for VMGC^50^ strain SGB020 and *p*=0.0027 overall, *p*=0.0014 within BMI≥25 for SGB182) and an unidentified *Bifidobacterium species* (*p*=0.0014 overall, *p*=0.0016 within BMI≥25; VMGC strain SGB065 **Fig. 2e-g**), which were all negatively associated with sPEC. Consistent with these associations, it has been previously shown that during pregnancy, gut *Bifidobacterium* levels are also negatively associated with the development of preeclampsia^54,55^.

We observed that the majority of taxa strongly associated with lower sPEC risk in this group belong to the species *Gardnerella vaginalis* (8 of 20, 0.001<*p*<0.012, 0.06<*q*<0.16; **Fig. 3c**), which has been frequently associated with bacterial vaginosis^56^, or to the genus *Bifidobacterium* (7 of 20, 0.002<*p*<0.017, 0.06<*q*<0.16; **Fig. 3c**), a Gram positive anaerobe that has various immunomodulatory effects^57,58^ and is the genus most closely related to *G. vaginalis*^59^. We next observed that 3 of the 4 taxa positively associated with the development of sPEC among individuals with higher BMI belong to the species *Prevotella timonensis* (0.005<*p*<0.016; 0.11<*q*<0.16). *Prevotella* species were previously shown to be associated with preterm birth^60^ and are major producers of sialidases^61,62^. Together, our results and these prior reports suggest that *G. vaginalis*, *Bifidobacterium*, and *P. timonensis* may play a role in the development of sPEC by regulating the inflammatory milieu of the reproductive tract. Examining vaginal immune factors also with respect to BMI, we have similarly found stronger and more numerous associations with sPEC in individuals with BMI≥25 (**Fig. S3**).

### Complex interactions among vaginal microbes, immune factors, and sPEC

The vaginal microbiome is associated with immune dysregulation and inflammation, both locally and systemically^63–66^, and we hypothesized that these interactions may be associated with the development of sPEC. We therefore investigated the relationships among these factors, starting with comparing the principal components of the vaginal immune factors with the robust principal components^51^ of the vaginal microbiome. We identified a significant correlation between microbiome RPC1 and the first two immune factor PCs (Pearson R=-0.38, *p*=4.5×10^−5^ and R=0.27, *p*=0.0038; **Figs. 4a**, **S4a**), as well as between immune factor PC1 and microbiome RPC3 (R=0.26 *p*=0.0026; **Fig. S4a**). These results support the hypothesis that the vaginal microbiome and local host immune profile in early pregnancy are partially associated, although there are independent components to each.

**Figure 4.**
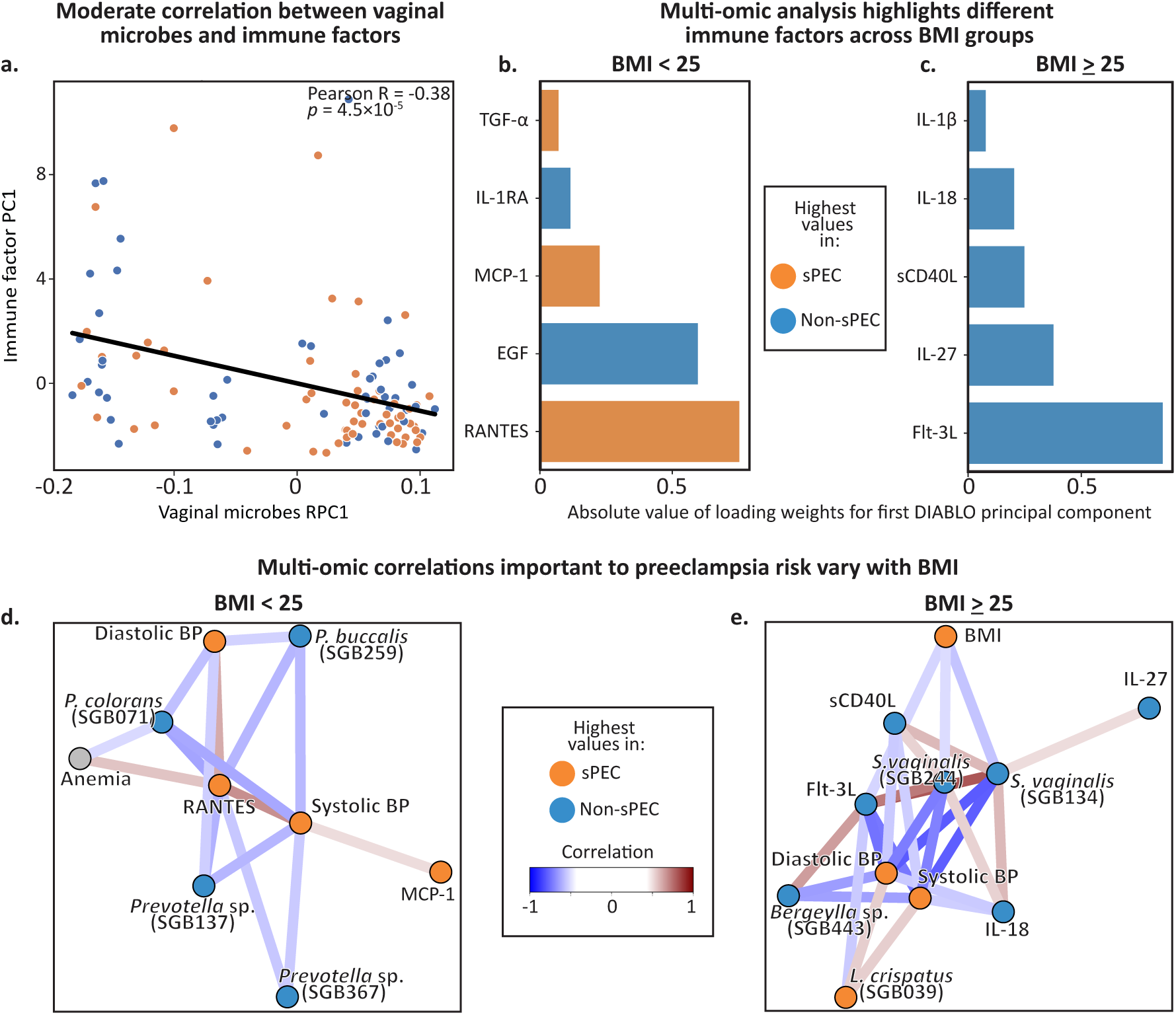
Complex interactions among microbiome, immune factors, and sPEC which vary with BMI. **a,** A scatter plot showing the first immune factors principal component (y-axis) and the first microbiome robust principal component (x-axis) for each individual in the cohort (N=110; Pearson R=0.39, *p*=2.4×10^−5^). **b,c** DIABLO loadings of the associations identified between immune factors and sPEC in a multi-omic analysis, within the BMI<25 (**b**) and BMI≥25 (**c**) groups. **d,e** Network of clinical, microbiome, and immune factor features identified via a DIABLO multi-omic analysis, restricted to a correlations cutoff of 0.5. Separate plots are shown for individuals with BMI<25 (**d**) and ≥25 (**e**).

To further investigate these patterns and identify interactions among vaginal microbes, immune factors, and sPEC, we ran a sparse discriminant analysis using DIABLO^67^, an integrative multi-omic analysis tool that identifies groups of features across multiple omics datasets that distinguish phenotypic groups^67^. Because we identified differences in associations of sPEC with vaginal microbes and immune factors between different BMI groups (**Figs. 3, S2, S3**), we performed this analysis stratified by maternal BMI. Even though DIABLO considers the associations between immune factors and sPEC in the context of how these features also relate to the vaginal microbiome and clinical characteristics, this analysis replicates several of our previous findings (**Figs. 2e-g**, **3e-g**). Consistent with our previous findings, the majority of immune factors were found to be reduced in individuals who developed sPEC (**Fig. 4b,c**, **S3b**). While most of these immune factors are secreted by immune cells, sCD40L is largely released by platelets^68^, which often become dysfunctional in preeclampsia, so its association with sPEC here may reflect subclinical platelet dysfunction early in pregnancy. DIABLO also identified three pro-inflammatory immune factors in individuals with BMI<25, TGFα, MCP-1, and RANTES, which were elevated in sPEC (**Fig. 4b**). MCP-1 and RANTES both recruit immune cells to sites of inflammation and to the uterus during pregnancy^69^, and were shown to be elevated the serum of individuals with PEC^70,71^. These higher levels in sPEC suggest that the vaginal ecosystem in individuals with BMI<25 may better reflect the systemic proinflammatory milieu that is characteristic of preeclampsia^72^.

To obtain better insights into the interactions among microbes, immune factors, and clinical features associated with sPEC, we next investigated the network of multivariate correlations inferred by DIABLO. In individuals with BMI<25, we found four *Prevotella* strains which were negatively associated with blood pressure (**Fig. 4d**). Conversely, while the role of vaginal *Prevotella* in hypertension is unknown, studies have suggested that high levels of *Prevotella* in the gut are associated with increased risk of developing hypertension^73,74^, suggesting an intriguing difference between *Prevotella* strains and the niche they occupy. We also found positive correlations between blood pressure and the proinflammatory immune factors MCP-1 and RANTES, which have previously been associated with hypertension and atherosclerosis^75,76^. We also observed that anemia was negatively correlated with a *P. colorans* strain and positively correlated with RANTES in this network (**Fig. 4d**). While anemia itself was not associated with sPEC (**Fig. S5**), its correlations with microbes and immune factors that are associated with sPEC may reflect the higher incidence of anemia during pregnancy in individuals with BMI<25^77^, which we also observed in our data (21% of individuals with BMI<25 had anemia, compared to 12% in BMI≥25). It may also suggest indirect mechanisms by which anemia may be involved in the early pathogenesis of sPEC.

In the DIABLO correlation network for individuals with BMI≥25, we identified several negative correlations between blood pressure and vaginal immune factors and microbes. Some of these associations, including a *Lactobacillus crispatus* strain that was positively associated with sPEC (albeit with a small coefficient in the DIABLO model; **Fig. S5e**) and blood pressure, and two *Sneathia vaginalis* strains that were negatively associated with sPEC and blood pressure (**Fig. 4e**), are surprising in light of the view of *L. crispatus* as a highly protective species^78–80^ and *S. vaginalis* as a pathobiont associated with chorioamnionitis and preterm birth^81,82^. We also identified several positive associations between *S. vaginalis* and immune factors that were lower in individuals who developed sPEC, including IL-18, IL-27, Flt-3L, and sCD40L. The positive associations between *S. vaginalis* strains and immune factors may reflect the highly immunogenic response that *S. vaginalis* can elicit during pregnancy^81^. We also identified strong correlations in this network between blood pressure and several immune factors, including Flt-3L, sCD40L, and IL-18. This similar involvement of multiple immune factors that otherwise belong to different signaling pathways may reflect a single underlying process with broad effects on vaginal immunity in sPEC. As before, the stark differences we observed between the associations identified in different BMI groups strongly suggest that underlying systemic inflammation and metabolic health may exert a strong influence on the vaginal ecosystem in sPEC. Altogether, our analysis reveals complex interactions between the vaginal microbiome, immune factors, and clinical factors that may reflect or play a role in the pathogenesis of sPEC.

### Vaginal immune factors and microbes predict sPEC months before diagnosis

Preeclampsia is typically not diagnosed until late in pregnancy^4^. We therefore investigated whether our clinical, vaginal microbiome or immune factor data, collected during the first trimester (mean±sd of 24.7±4.6 weeks prior to sPEC diagnosis), hold potential for improving early sPEC diagnosis. We trained logistic regression models and evaluated their performance on held-out samples using nested rebalanced^83^ leave-one-out cross validation, such that model hyperparameters were optimized using internal cross-validation within each training set, and a model with these chosen parameters was then evaluated once on the held-out test sample, without data leakage (**Fig. S6; Methods**). Given the interactions we observed with BMI for both microbiome and immune associations with sPEC, BMI stratification was implemented as a parameter that our models could utilize (**Methods**).

As a benchmark, we first devised models predicting sPEC using only clinical information, such as pre-pregnancy history of blood clots, hypertension, and heart disease (**Methods**). These models obtained reasonable accuracy (auROC=0.62, area under the precision-recall curve [auPR]=0.64, Mann-Whitney *U* test vs. a null of random prediction *p*=0.032, **Methods**; **Fig. 5a,b**), on par with previous research^84–86^. Training similar models using microbiome data obtained a stronger performance (auROC=0.71, auPR=0.67, *p*=1.1×10^−4^; **Fig. 5a,b**), and similar results were obtained using immune factor levels (auROC=0.72, auPR=0.73, *p*=7.9×10^−5^; **Fig. 5a,b**). We observed that 99% of microbiome-based models that were ultimately selected by cross validation within the training data (**Methods**) utilized BMI stratification, compared to only 2% of immune factor-based models. These predictive associations, which generalized to held-out samples with conservative evaluation, further demonstrate the robust association between the development of sPEC and the early pregnancy vaginal microbiome and immunity.

**Figure 5.**
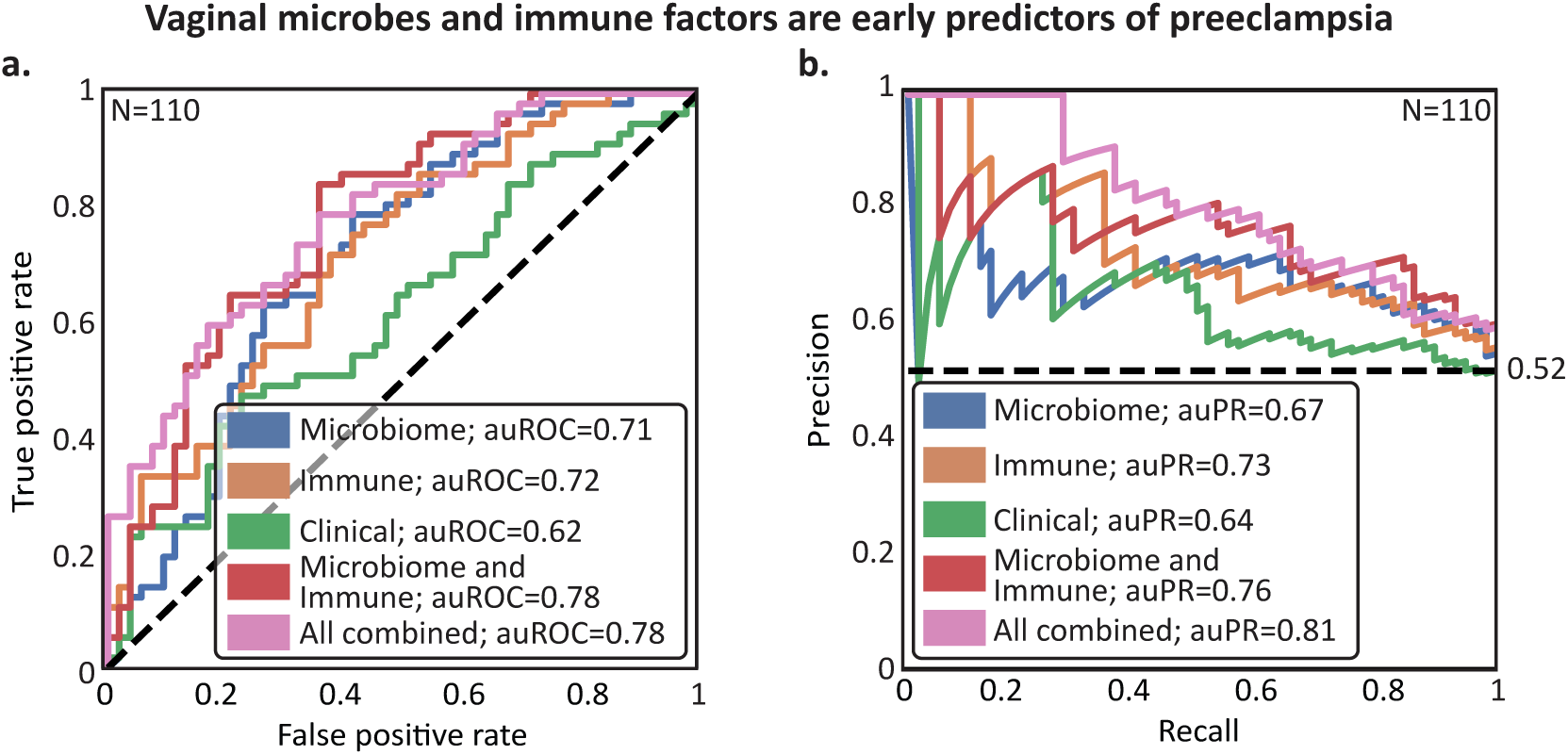
The early pregnancy vaginal microbiome and host immunity are predictive of sPEC ~6 months in advance. **a,b** Receiver Operating Characteristic (**a**) and Precision-Recall curves (**b**) of leave-one-out nested cross-validation predictions of linear models (**Methods**). Dashed horizontal line in (b), class balance. Results are shown for models considering only the microbiome, immune factor, and clinical data, respectively, along with combinations of the microbiome and luminex predictions, and those of all three models.

When combining the predictions of the different modalities, we obtained additional improvements. Combining microbiome and immune factor data achieved an auROC of 0.78 and auPR of 0.76 (Mann-Whitney *U p*=5.7×10^−7^, **Methods**; **Fig. 5a,b**). Further incorporating clinical data improved results slightly (auROC=0.78, auPR=0.81, *p*=2.7×10^−7^; **Fig. 5a,b**). The performance of our multi-modal models illustrates that microbiome and immune factors each add independent information concerning sPEC risk, as opposed to clinical data (other than BMI) which does not substantially improve prediction accuracy. Despite being evaluated in a conservative nested cross-validation framework, the final performance of our model, with an auROC of 0.78, is superior to models based on blood proteins^87^. While this performance is comparable to models based on other molecular data, such as serum cell-free RNA^9^ or PlGF and sFlt-1^8^, which each achieve an auROC of 0.82, our models notably use data collected substantially earlier in pregnancy (first trimester compared to second trimester for cell-free RNA and late pregnancy for PlGF and sFlt-1).

### sPEC prediction models are accurate in an independent dataset

To further demonstrate the potential of vaginal microbiome and immune factor data for sPEC diagnosis early in pregnancy, we next tested if models trained on data from our cohort can generalize to data from an independent external cohort. To this end, we used data from the MOMS-PI study^22^, which generated metagenomic sequencing and immune factor data from samples collected early in pregnancy. 33 participants with available early pregnancy data in this cohort self-reported whether they developed eclampsia or preeclampsia. While multiple samples were collected per participant in the MOMS-PI cohort, we used the earliest sample collected before week 17 of gestation. We processed metagenomic sequencing data from this cohort using the same pipeline we used to process data from the nuMoM2b cohort, retaining 17 individuals with >500,000 non-human reads (**Methods**), with 5 of them reporting a positive diagnosis. Of note, these participants differed significantly from our nuMoM2b cohort in their BMI (Mann-Whitney *U p*=0.0091) and parity (*p*=7.2×10^−7^), two factors known to impact the vaginal ecosystem^52,88^, as well as in self-identified race (*p*=3.1×10^−6^; **Table 2**). Additionally, only 11 of the 20 immune factors measured in nuMoM2b were available for this cohort, and the only clinical features available were maternal age, blood pressure, BMI, and weight. We corrected processing bias in the microbiome data with DEBIAS-M^89^, to which we applied models that were trained and selected using cross-validation within the nuMoM2b cohort to the MOMS-PI dataset without retraining (**Fig. 6**).

**Table 2.** MOMS-PI subcohort characteristics. GA, gestational age; *p*, Mann-Whitney *U* or Fisher’s Exact, as appropriate. Age was missing for one MOMS-PI participant.

| | nuMoM2b | MOMS-PI | Difference<br>($p$ value) |
| --- | --- | --- | --- |
| **N** | 124<br>[62 sPEC] | 17<br>[5 PEC or<br>eclampsia] |  |
| **Race [N (%)]** |  |  | **$3.1\times 10^{-6}$** |
| **Non-Hispanic Black** | 30 (24.2) | 16 (94.1) |  |
| **Non-Hispanic White** | 52 (41.9) | 1 (5.9) |  |
| **Other** | 42 (33.9) | 0 |  |
| **BMI (median kg/m<sup>2</sup> [IQR])** | 26.3<br>[22.9-32.2] | 34.4<br>[28.1-38.9] | **0.0091** |
| **N total pregnancies**<br>**(median [range])** | 1<br>[1-5] | 3<br>[1-9] | **$7.2\times 10^{-7}$** |
| **GA at delivery**<br>**(median weeks [range])** | 38.8<br>[21.1-42.1] | 38.4<br>[25.6-41.1] | 0.30 |
| **Age years (N [%])** |  |  | 0.99 |
| **<18** | 4 (3.2) | 0 |  |
| **18-29** | 76 (61.3) | 11 (64.7) |  |
| **29-38** | 39 (31.5) | 5 (29.4) |  |
| **>38** | 5 (4.0) | 0 |  |
| **GA at sample collection**<br>**(median weeks [range])** | 12<br>[6-14] | 11<br>[6-15] | **0.021** |

**Figure 6.**
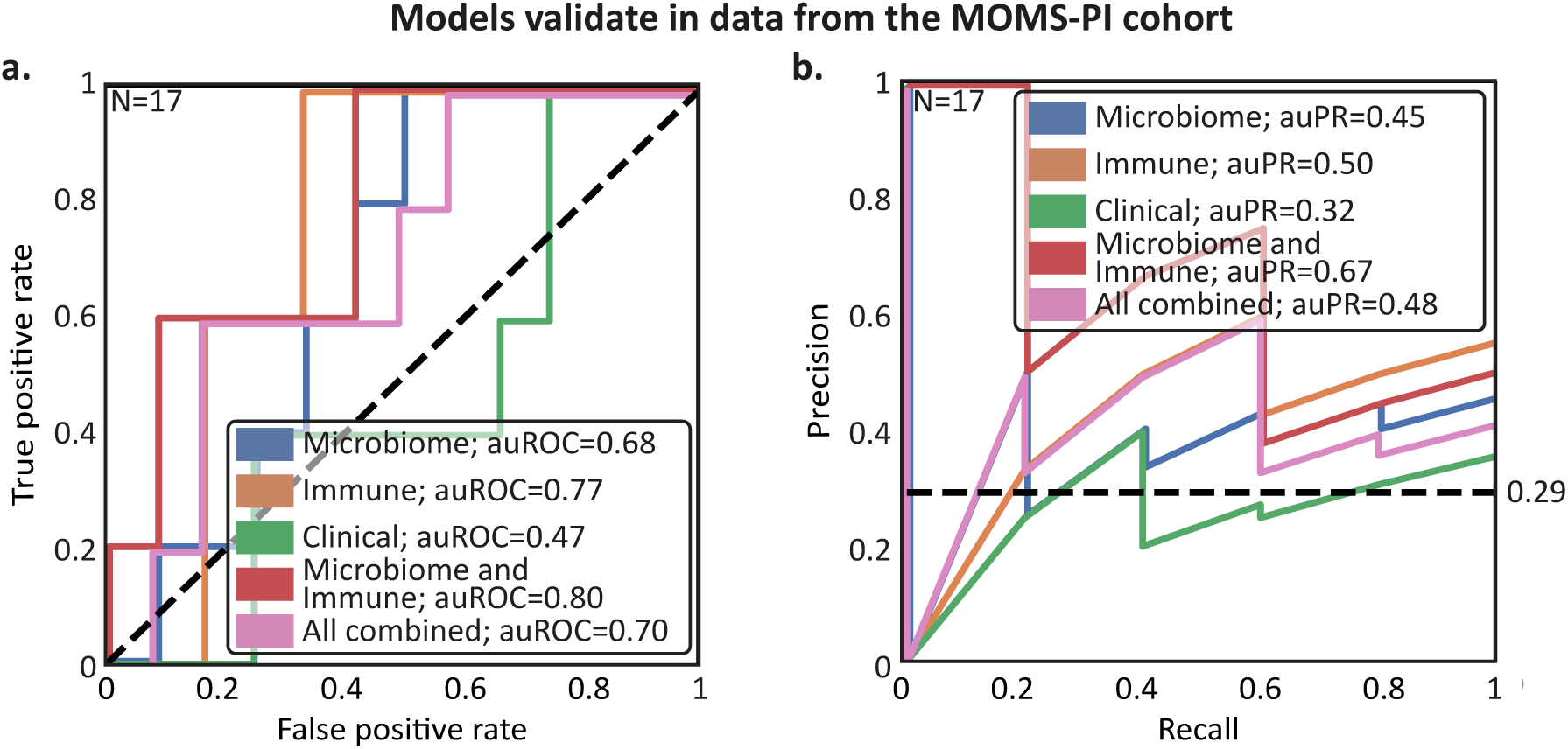
Predictions of sPEC ~6 months in advance using early pregnancy vaginal microbiome and host immunity generalize to independent cohorts. **a,b** Similar to **Fig. 5a,b**, but for models trained on data from the nuMoM2b cohort and tested on data from the Multi-Omic Microbiome Study-Pregnancy Initiative (MOMS-PI) collected by Fettweis et al^22^.

This is a challenging benchmark, which includes generalization to a new population with different characteristics (**Table 2**), an independent study with different methods and investigators, and differences in both measurement techniques and outcome assessment. This is underlined by our finding that models based on clinical data did not generalize well between the studies (auROC=0.47, auPR=0.32, **Fig. 6a,b**). Nevertheless, we saw similar predictive performance and successful generalization for the microbiome (auROC=0.68, auPR=0.45), and immune factor-based models (auROC=0.77, auPR=0.50; **Fig. 6a,b**). We also observed similar performance and successful generalization when we evaluated the combined models, as our model based on microbiome and immune factor data achieved an auROC of 0.80 (auPR=0.67, **Fig. 6a,b**). These results, demonstrating that the association between preeclampsia and the early-pregnancy vaginal microbiome and host immunity generalizes across cohorts, underscore the robustness of our analysis and the potential of these measurements to improve early diagnosis of sPEC.

## Discussion

In this study, we quantified the levels of microbes and immune factors in the vaginal ecosystem using samples collected early in pregnancy (first trimester) from 124 pregnant individuals. We found significant associations between both vaginal microbes and immune factors and the development of sPEC, which differ with maternal BMI, and involve known preeclampsia risk factors such as blood pressure and anemia. Integrating immune factor, microbiome and clinical data allows for prediction of sPEC with comparable accuracy to previous molecular tests^8,90^, but much earlier in pregnancy. Finally, we demonstrated the robustness and generalizability of the associations we detected by validating our predictive models in an independent external cohort^22^.

We identified several immune factors significantly reduced in the vaginal ecosystem of individuals who developed sPEC, including IL-1RA, Flt-3L, and EGF. These associations may be due to compromised immune function within the vaginal ecosystem and perhaps along the entire reproductive tract, as it is known that immune cells are integral to maintaining the protective vaginal epithelium^91^ and play a role in spiral artery remodeling^92^. This is echoed in recent investigations of placentas of women with preeclampsia, which exhibit reduced numbers of macrophages during delivery, and lower local expression of innate immune factors^93^. The reduction in placental immune cells is also associated with decreased spiral artery remodeling in mice^94^. At the same time, some of the vaginal immune factors that we found to be reduced in individuals who developed sPEC have been positively associated with preeclampsia when measured in serum late in pregnancy^48,49^. This discrepancy suggests that immune factor levels in the local vaginal environment may reflect biological processes that do not manifest in systemic circulation, and motivate additional research to understand the interplay between local and systemic immunity in preeclampsia.

Our results highlight a potential subgroup effect, which manifested as a strong modification by BMI of the relationship between sPEC development, the vaginal microbiome, and immune factors. Our results echo those of another study, which showed that the association between adverse pregnancy outcomes and the serum sFlt-1/PIGF ratio, a well-studied preeclampsia risk factor^8^, is significantly weaker in obese individuals^95^. Differences in BMI are associated with metabolic health and systemic inflammation, which would likely be related to dysregulation of microbes or immune factors in the reproductive tract. As obesity dysregulates many biological pathways, including estrogen, progesterone^96,97^, and vaginal glycogen metabolism^98^, our findings motivate mechanistic studies to determine the directionality and specific mechanisms that may underlie the interaction of BMI with the associations of vaginal microbes and immune factors in sPEC that we detected here.

We found several intriguing associations between vaginal microbes and sPEC in individuals with BMI≥25. These included several strong associations between sPEC and *P. timonensis,* a major source of sialidases in the vaginal ecosystem, which can degrade the mucin lining of the reproductive tract and elevate the risk of preterm birth^99,100^. A previous study has also found that another *Prevotella* species, *P. bivia*, was enriched at the time of birth in individuals with preeclampsia^101^. We also found that several *Bifidobacteria* strains were negatively associated with sPEC. *Bifidobacteria* can be highly abundant in the vaginal microbiomes of healthy reproductive-aged individuals, and were shown to produce lactic acid at comparable levels to *L. crispatus*^102^, which promotes a healthy vaginal pH and epithelial barrier function^103^. Additional investigations could determine if elevated sialidase activity or lactic acid levels underlie these associations with sPEC.

In individuals with BMI≥25 we also found that several *G. vaginalis* and *S. vaginalis* strains were negatively associated with sPEC, a surprising result given that these taxa are often associated with dysbiosis and adverse outcomes^22,81,82,104,105^. We note, however, that these taxa are often part of diverse vaginal microbiomes, and that our results may indicate that within such diverse ecosystems, low levels of these taxa, and thus higher levels of other taxa such as *Fannyhessea vaginae*, are associated with preeclampsia. Alternatively, the protective association of *S. vaginalis* may be related to its immunogenic capacity during pregnancy^81^, contrasting the reduced levels of immune factors we found in individuals who developed sPEC. Additional mechanistic studies are needed to elucidate the role of these microbes in the reproductive tract during pregnancy and their interplay with the immune system to determine if they are causally implicated in the pathogenesis of sPEC.

Our multi-omic analysis has also identified several associations between vaginal immune factors and other clinical features, such as blood pressure and anemia. These include positive correlations between blood pressure in individuals with BMI<25 and the proinflammatory cytokines MCP-1 and RANTES, which are known to contribute to hypertension and atherosclerosis^75,76^, as well as an association between RANTES and anemia, which while not associated with sPEC in our cohort, has previously been reported to increase the risk of developing preeclampsia^106^ and other adverse pregnancy outcomes^107^. We also identified negative associations between blood pressures and IL-18, sCD40K and Flt-3L in individuals with BMI≥25. These associations are discordant with positive associations with hypertension that were previously identified when these factors were measured in blood^108–110^, once again highlighting a difference between local and systemic immune factor measurements.

Our predictive modeling approach has several limitations. First, our use of a case-control cohort enriched for sPEC limits our ability to assess population-level predictive value, and further validation is required in prospective studies. Second, while our predictive models demonstrate accuracy in two independent cohorts, both are limited to individuals residing in the United States. Additional studies in global populations are needed to further evaluate generalizability and identify robust preeclampsia biomarkers. Third, while our use of BMI allowed us to identify a significant interaction affecting the relationships among vaginal microbes, immune factors, and sPEC development, BMI alone provides an incomplete picture of metabolism^111^. To better capture metabolic health, future studies may benefit, for example, from incorporating serum lipid or blood glucose levels. Finally, integration of other data types, such as cell free RNA measurements or sFlt-1 and PlGF, which have shown some promise in preeclampsia prediction^8,90^, may facilitate even stronger predictive signals.

Currently, preeclampsia is diagnosed based on symptoms indicating impending multi-organ damage that manifests late in pregnancy. Despite its high prevalence and severe health consequences for both mother and child, we lack effective methods for early identification of individuals in which the pathogenesis of preeclampsia is already taking place without overt symptoms. The lack of success in identifying biomarkers of preeclampsia, including in a large investigation of plasma proteomics^112^, further underscores the difficulty inherent in the study of preeclampsia pathogenesis. Improved methods to diagnose preeclampsia during early pregnancy would offer an opportunity to dramatically improve outcomes through increased monitoring and earlier interventions. Here, we identified a generalizable signal within the first trimester vaginal ecosystem that predicts the future diagnosis of sPEC with comparable accuracy to other tests that are currently being developed^8,9^, which use data collected significantly closer to the typical time of diagnosis. Altogether, our results demonstrate that investigating the role of the early pregnancy vaginal ecosystem in preeclampsia may provide an improved understanding of preeclampsia pathogenesis and promising leads for identifying prevention and treatment strategies.

## Methods

### Study design and cohort description

We analyzed samples that were collected as part of the multi-center nuMoM2b study^29^. This study enrolled nulliparous pregnant individuals at the following clinical sites: Case Western University; Columbia University; Indiana University; University of Pittsburgh; Northwestern University; University of California Irvine; University of Pennsylvania; and University of Utah, and was approved by the Institutional Review Board at each participating clinical site. The current analysis was approved by the IRB of Columbia University, approval number AAAT7071.

Between 2010 and 2015, the nuMoM2b study enrolled 10,038 pregnant participants between 6 and 14 weeks of gestation. Pregnancy dating was confirmed via ultrasound crown-rump length measurement. Study participants answered questionnaires and provided biospecimens during a visit between 6 and 14 weeks of gestation, in which 2 cervicovaginal swabs were used by each study participant to self-collect cervicovaginal fluid. One swab, intended for analysis of vaginal proteins and metabolites, was suspended in 2 mL cryovial w/1 mL DPBS, and the other swab, intended for analysis of microbial DNA, was suspended in a 2 mL cryovial with 1 mL 10 mM Tris-Hcl, 0.1 mM EDTA.

### Chart abstraction and preeclampsia definition

Chart abstractors used data collected at each study visit as well as prenatal records, records from antepartum hospitalizations, and records from labor and delivery to determine the baseline hypertension and proteinuria status of each study participant and determine whether they developed any hypertensive disorders during their pregnancy. The nuMoM2b study employed multiple definitions for preeclampsia with and without severe features, one of which was based on the ACOG 2013 guidelines^113^, which was the definition used for this study. New onset gestational hypertension was defined as systolic blood pressure of 140 mmHg or greater or diastolic blood pressure of 90 mmHg or greater on two occasions six hours apart or on one occasion that then prompted antihypertensive treatment. Preeclampsia with severe features (sPEC) was defined as new onset gestational hypertension plus any of the following symptoms: thrombocytopenia (platelet count < 100,000/*µ*l), pulmonary edema, serum creatinine >1.1 mg/dL, severe headache, scotoma, serum aspartate aminotransferase (AST) >100 IU/L, epigastric pain, or severe hypertension (systolic blood pressure≥160 or diastolic blood pressure≥110 on 2 occasions ≥6 hours apart or on 1 occasion requiring antihypertensive therapy and excluding blood pressure readings during the second stage of labor).

### Identifying clinical features for association and prediction

nuMoM2b study participants provided extensive sociodemographic and clinical data at a screening visit that occurred between 6 and 14 weeks of pregnancy. To construct a dataset of clinical features for multi-omic association analysis as well as for prediction, we relied on previous work that used clinical data to predict sPEC in the nuMoM2b cohort^86^ as well as ACOG guidelines regarding PEC risk factors^114^. We used the following variables in our multi-omic association analysis with DIABLO: history of liver disease, gallbladder disease, diabetes prior to pregnancy, autoimmune disease, endocrine disease, blood clots or thromboembolic disease, heart disease, kidney disease, anemia, or hypertension, as well as first trimester blood pressure readings, maternal age, maternal BMI, maternal education level and maternal socioeconomic status. For predictive modeling we used the following variables: history of liver disease, gallbladder disease, diabetes prior to pregnancy, autoimmune disease, endocrine disease, blood clots or thromboembolic disease, heart disease, kidney disease, anemia, or hypertension, as well as first trimester blood pressure readings, maternal age, and maternal BMI. All clinical data was ascertained or measured at first trimester visits.

### Subject and sample selection for immune factor and microbiome profiling

62 individuals who developed sPEC were randomly chosen for immune factor and vaginal microbiome profiling. These 62 cases were matched with 62 participants who did not develop sPEC, eclampsia, or superimposed preeclampsia, during their pregnancy, although these individuals may have developed gestational hypertension or mild preeclampsia. These individuals were frequency matched based on a combination of maternal age and self-identified race, as well as enrollment site.

### Immune factor profiling

The levels of the following immune factors were quantified in 150 ul aliquots of the samples intended for protein and metabolite analysis via a multiplexed Luminex assay (Luminex Corporation) using Millipore kits (HCYTA-60K-PX48 and HCYTA-60K): EGF, Flt-3L, G-CSF, IFNα2, IFN-γ, IL-10, IL-18, IL-1RA, IL-1α, IL-1β, IL-27, IL-4, IL-6, IL-8, MCP-1, RANTES, TGFα, TNFα, VEGF, and sCD40L. All measurements were obtained in duplicate and data were in units of pg/ml. Immune factor profiles for the 124 participants were obtained on two separate days, with 84 samples profiled (42 sPEC cases, 42 controls) on one day, and 40 samples (20 sPEC cases, 20 controls) on another. There was no confounding association between processing batch and sPEC (Fisher’s Exact *p*=1.0). To account for technical variation between batches, data was batch-standardized by subtracting the mean and dividing by the standard deviation of each feature in each batch. Concentrations and limits of detection were determined following the construction of a standard curve for each analyte. Values below the limit of detection were imputed with the limit of detection specific to each immune factor of the corresponding batch, and divided by √2. Values that were obtained with a high coefficient of variation (>20%) were replaced and imputed with the mean value of the corresponding batch for each immune factor.

### Microbiome profiling

Microbial DNA was extracted from 200 ul aliquots of the sample collected for microbial DNA analysis using the QIAcube HT and either the QIAmp 96 Virus QIAcube HT kit (Cat. No. 57731) or the Qiagen DNeasy PowerSoil HTP 96 kit (Cat. No. 12955-4-5D). The former kit was used to extract DNA from 27 sPEC cases and 27 controls, while the latter kit was used to extract DNA from all other samples. Extracted DNA was then prepared for sequencing via the Nextera DNA Library Prep Kit (Cat. No. 20060059). Metagenomic sequencing data was generated for each sample via paired end sequencing (2×100bp) using a NovaSeq 6000 to a mean±std depth of 43.23 ± 23.4 million reads. Samples were processed in batches, which were explicitly balanced with respect to sPEC, clinical enrollment site, small for gestational age, maternal BMI, maternal age, maternal self-identified race, smoking status, maternal socioeconomic status, spontaneity of birth, chorioamnionitis, short cervix, cesarean birth, fetal sex, hypertensive disorders, live birth, gestational age, and prelabor membrane rupture, to ensure a lack of batch confounding. Therefore there was no association between study group and kit or batch status in our experimental design (Fisher’s exact *p*=1.0 for kit, *p*=0.99 for batch).

Metagenomic sequencing reads from each sample were quality filtered using Trimmomatic^115^ (v0.39) in order to remove reads containing Illumina adapter sequences, bases with quality scores below 25, and reads shorter than 50 bases. Metagenomic reads were then aligned to the human genome and PhiX genome via Bowtie2^116^. Any reads where either end mapped were subsequently removed^117^. Following the removal of host DNA, all samples were subsampled to a consistent depth of 500,000 microbial reads to normalize the data across our dataset. Any sample with less than 500,000 reads was discarded from further analysis, leaving 110 samples (N sPEC = 57). Microbial abundances were then estimated using Kraken2 (v2.1.3)^118^ followed by Bracken (v2.9)^119^. In order to ensure the most accurate and relevant results, we utilized the Human Vaginal Microbiome Genome Collection (VMGC)^50^ as the reference database, which is specifically curated to include a comprehensive and diverse array of vaginal microbial genomes. As the VMGC classified *G. vaginalis*, *G. piotii*, *G. leopoldii*, and *G. swidsinskii* as species within the genus *Bifidobacterium*, we reclassified these species as members of the *Gardnerella* genus. All unmapped reads were accounted for in the final read count matrix in an ‘unmapped’ column. For all analyses, the microbiome data was preprocessed by converting to relative abundances followed by a center log transform with a pseudocount of 10 to the power of the smallest whole number that keeps the pseudocount below the smallest observed relative abundance value across the dataset.

### Association of immune factors and microbes with sPEC

To evaluate associations between the microbiome data and sPEC, we performed two-sided Mann-Whitney *U* tests comparing the relative abundance of each microbe with a minimum of 10 detected reads in a minimum of 10 samples per group between the sPEC and non-sPEC groups, among samples within which a given microbe was detected. Additionally, we performed a robust principal component analysis^51^ to compare the entire microbial communities between the two groups. We investigated the relationships among microbes, immune factors, sPEC, and BMI via DIABLO^67^. We performed the analysis for 1) the entire cohort; 2) the 43 individuals with a BMI<25; and 3) the 67 individuals with a BMI≥25. Because nonrestrictive presence thresholds produced a high stochastic variation in results, we ensured consistency for the DIABLO analysis by reducing the number of features; only the microbes present in abundance of 10^−3^ in a minimum of 5% of samples were considered. The relevance network for the fitted DIABLO models were plotted using a ‘cutoff’ parameter of 0.5.

### Predicting risk of preeclampsia

To determine whether first trimester immune factor, microbiome, and clinical data is predictive of later diagnosis of sPEC, we trained and evaluated models using L^2^ regularized logistic regression. All predictive model pipelines were evaluated using a rebalanced leave-one-out cross-validation scheme^83^, with an inner (nested) 5-fold cross-validation to tune hyperparameters **(Fig. S6**). The hyperparameters tuned in this analysis were the number of features to select via ANOVA F-value ([2, 5, 10]), implemented via scikit-learn’s^120^ ‘SelectKBest’, and the L^2^ regularization strength in the linear model which followed scikit-learn’s default cross-validation range from 1 to 10^8^. The nested tuning also considered models stratified by BMI above and below 25, which, like other hyperparameters, was performed only on the inner folds. To apply the rebalanced leave-one-out approach with stratification for BMI, the additional held-out point^83^ was randomly selected from the same BMI group as the held-out test point. Combined models were constructed by averaging the predictions of each individual predictor across the three data types – microbiome, immune factor, and clinical. To determine the statistical significance of our models’ auROCs vs. a random null, we used the Mann-Whitney *U* test to compare the predictions made for sPEC and non-sPEC^121^.

### Assessing generalizability of microbiome, immune factor, and clinical models

To assess the ability of our models to generalize to data from the Multi-Omic Microbiome Study-Pregnancy Initiative (MOMS-PI) cohort^22^, raw microbial reads and immune factor measurements were downloaded from dbGaP, study no. 20280 (accession ID phs001523.v1.p1). MOMS-PI samples were included from study participants who responded either yes or no during their delivery visit as to whether they experienced preeclampsia or eclampsia during pregnancy. Metagenomics data was processed as described above, and microbial abundances were also estimated using the VMGC reference database^50^. Prior to validation, the two microbiome datasets were preprocessed using log-additive DEBIAS-M^89^ with default parameters, while only observing metadata from the nuMoM2b samples. Models based on clinical and immune factors data were trained and evaluated using only features shared by both cohorts. For the clinical models, this included only maternal age, BMI, weight, systolic blood pressure, diastolic blood pressure, and mean arterial pressure. For the immune factors, this included only 11 immune factors: IL-8, IL-4, IL-10, RANTES, TNFα, IL-6, IL-1β, IL-1RA, MCP-1, G-CSF, and IFNγ. A single validation model was selected from within nuMoM by Rebalanced Leave-One-Out cross-validation to tune the same set of hyperparameters listed above. Prior to validation, the immune factor and clinical measurements from the MOMS-PI cohort were standardized using scikit-learn’s StandardScaler class. When combining multiple models together, we first standardized each model’s predictions using StandardScaler.

## Acknowledgements

We thank members of the Korem group, the nuMoM2b study consortium, and the nuMoM2b-HHS2 study consortium for useful discussions. We thank all participants, staff, and investigators involved in the generation of data used in this study. This study was supported by grant funding from the Eunice Kennedy Shriver National Institute of Child Health and Human Development (NICHD): R01 HD106017 (T.K.), R01 HD114715 (T.K.), 5F30 HD108886 (W.F.K.), U10 HD063036, U10 HD063072, U10 HD063047, U10 HD063037, U10 HD063041, U10 HD063020, U10 HD063046, U10 HD063048, and U10 HD063053; the National Library of Medicine: T15 LM007079 (G.I.A, J.A.U.); the National Institute of General Medical Sciences: T32 GM007367 (W.F.K); the National Heart, Lung, and Blood Institute: U01 HL145358; the National Cancer Institute: R01 CA245894 (A.-C.U.) and P30 CA013696 (for the Genomics and High Throughput Screening Shared Resource); and the Clinical and Translational Science Institutes: UL1 TR001108 and UL1 TR000153. Data from the MOMS-PI dataset was obtained from dbGaP (phs001523), and additional data was provided by Gregory A. Buck, Ph.D. and colleagues and supported by NICHD (U54 HD080784) (G.A.B.).

## Author contributions

W.F.K, A.B.K and T.K. designed the study. W.F.K. and G.I.A. designed and conducted all analysis, interpreted the results, and contributed equally to this work. Y.M., J.A.U., and A.B.K assisted with data analysis. H.P. and E.W. processed samples and generated microbiome data, under the supervision of A-C. U. S.P. processed samples and generated immune factor data, under the supervision of R.N. M.S. and G.A.B. assisted with data access and acquisition. W.F.K., G.I.A., Y.M. and T.K. wrote the manuscript with critical input from all authors. T.K. and R.J.W. conceived the study. T.K. supervised the study.

## Supplementary Figures

**Figure S1.**
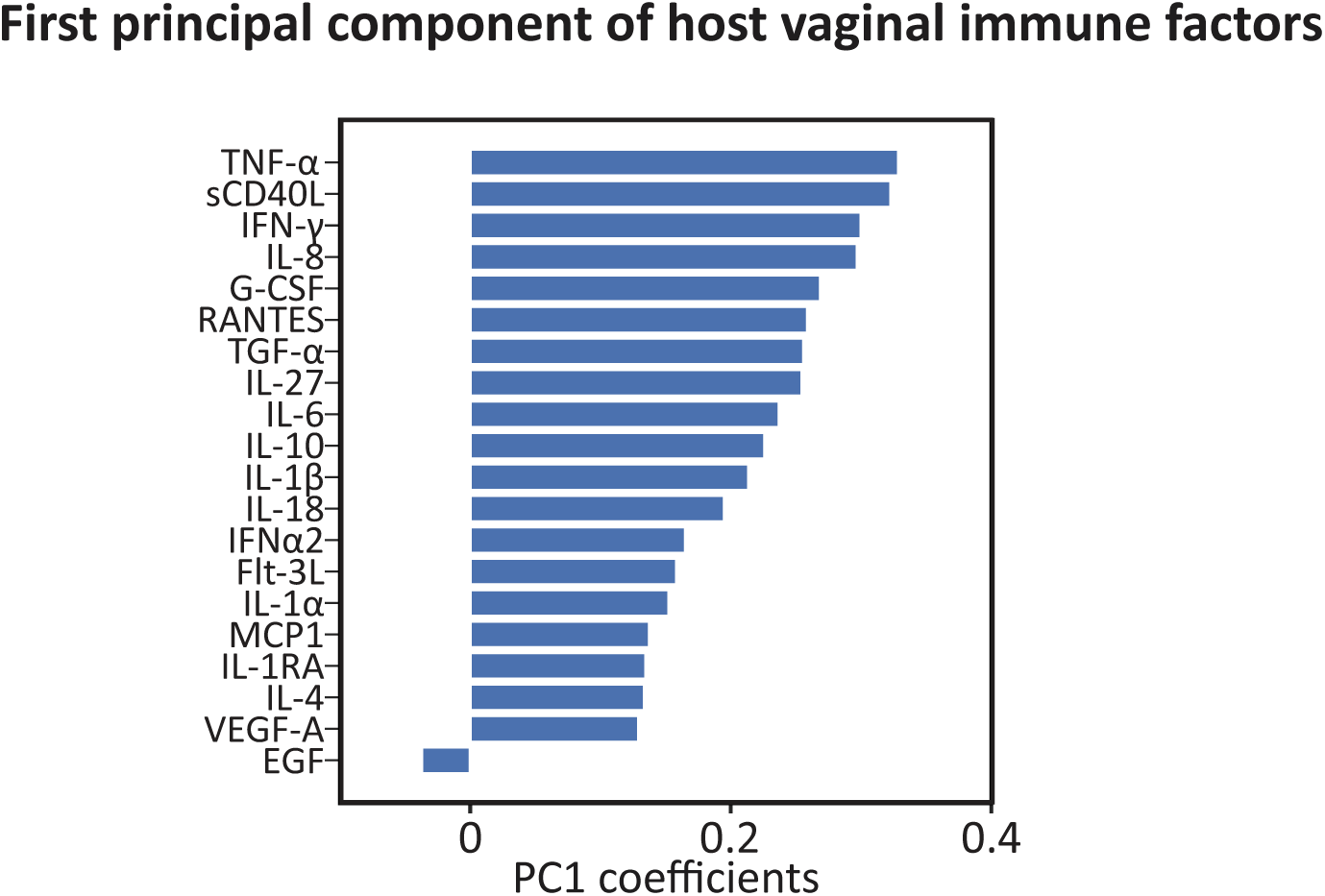
The association of immune factor PC1 with sPEC is driven by multiple immune factors. Barplot of the coefficients of the immune factors PC1.

**Figure S2.**
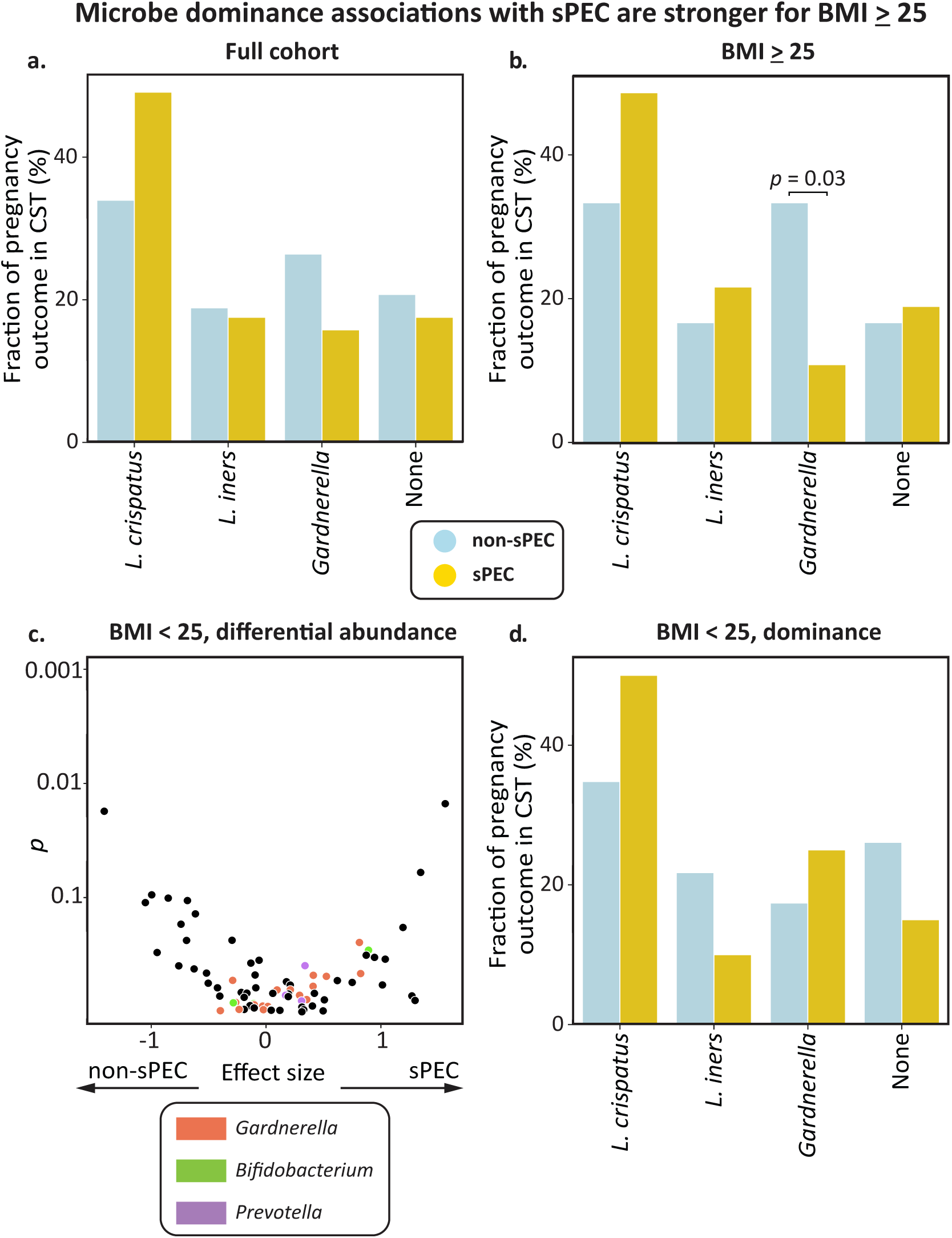
Weak associations between vaginal microbes and sPEC in dominance analysis and in individuals with BMI<25. **a,b,** Bar plots showing the fraction of individuals with and without sPEC, separated by vaginal microbiome dominance. Dominance was defined as >30% relative abundance, following previous studies^22^. None, no microbiome with >30% abundance; *p*, Fisher’s exact. **c,** Similar to Fig. 3c,d, but for individuals with first trimester BMI<25. **d,** Same as (b) for individuals with first trimester BMI<25.

**Figure S3.**
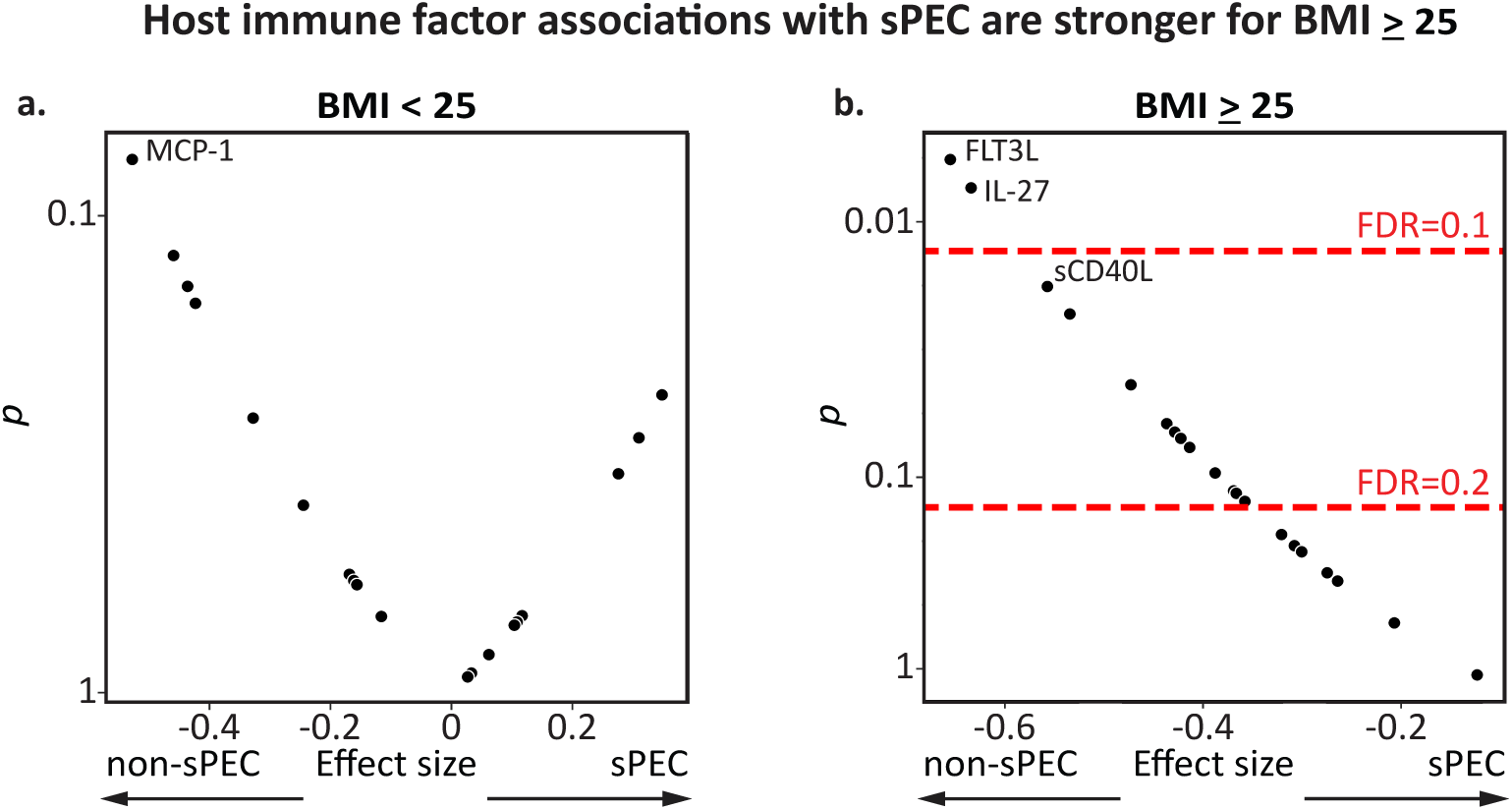
Vaginal immune factors are more strongly associated with sPEC in individuals with BMI≥25. **a,b,** Same as Fig. 2d, stratified by BMI.

**Figure S4.**
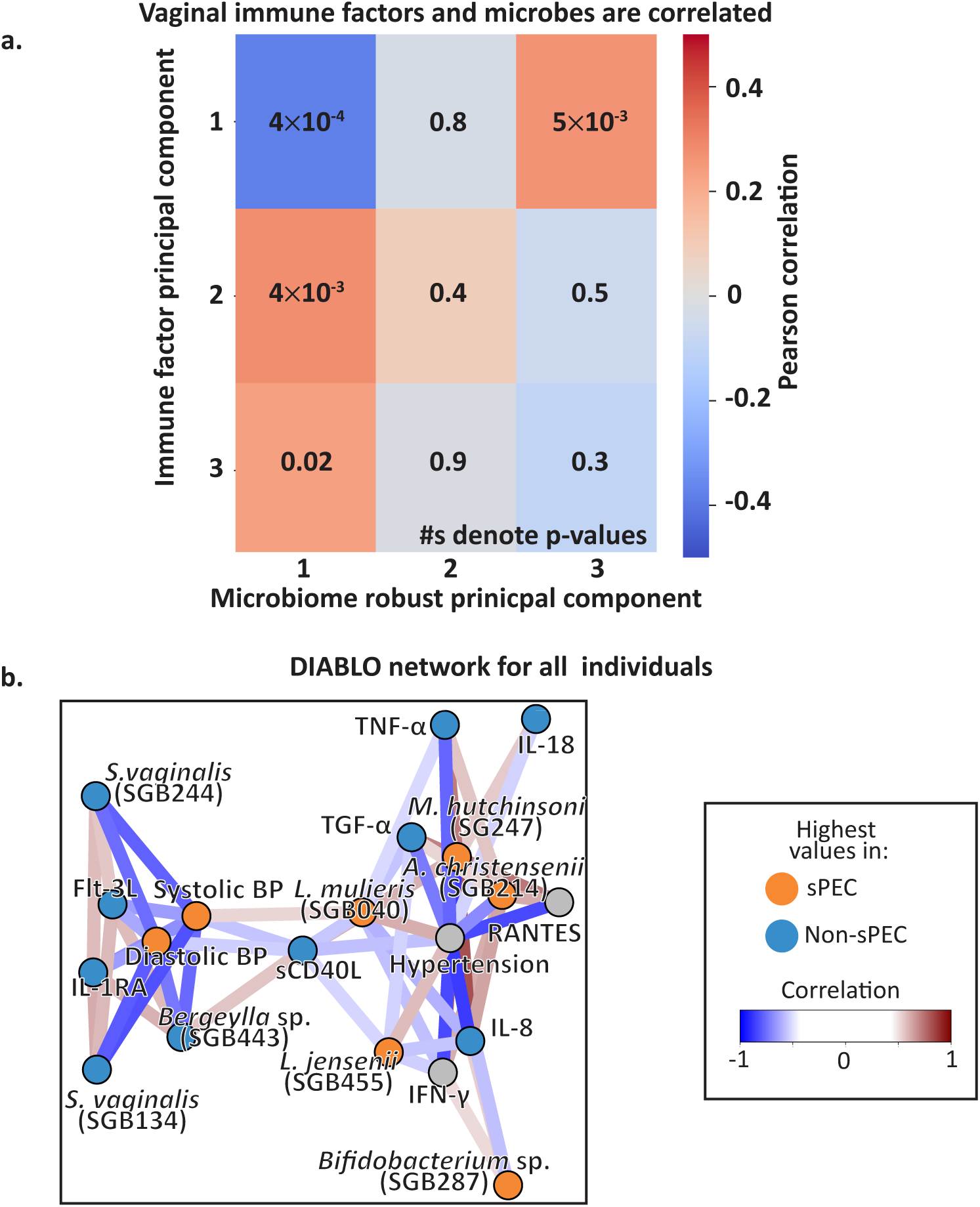
Associations between vaginal microbiome and immune factors in the context of sPEC. **a.** Pearson correlations of the first three microbiome RPCs and the first three immune factors PCs. **b.** Similar to Fig. 4d,e, but for a DIABLO analysis which included all individuals regardless of BMI.

**Figure S5.**
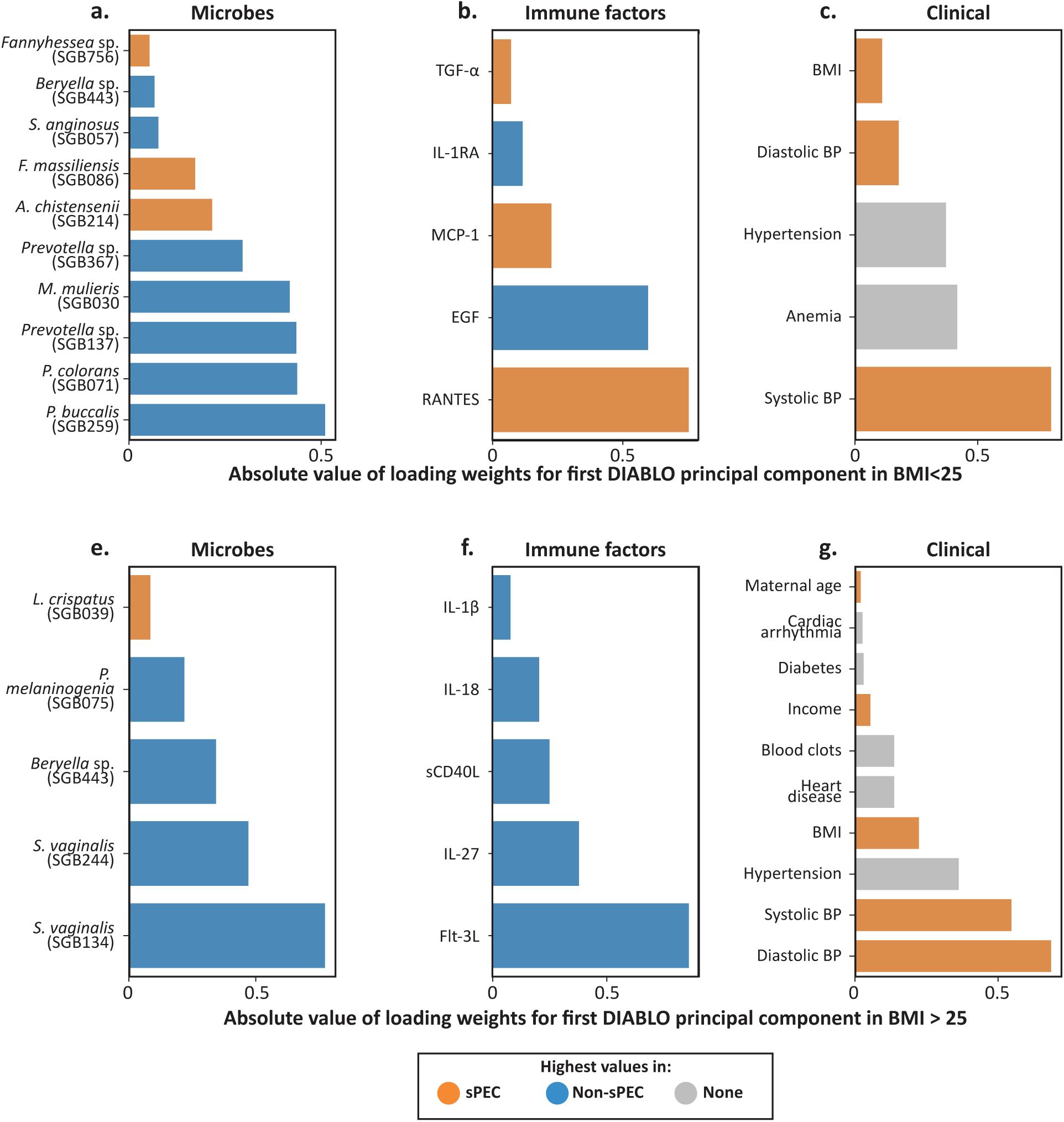
Associations of microbes, immune factors, and clinical covariates with sPEC in a sparse multivariate model. **a-c,** Coefficients of fitted DIABLO model’s associations with the first principal component of each data modality when trained on individuals with BMI<25, for microbes (**a**), immune factors (**b**), and clinical data (**c**). Colors of the bar denote the sPEC status that each variable is associated with. **d-f**, Same as a-c, but for a DIABLO model fitted using data from individuals with BMI≥25.

**Figure S6.**
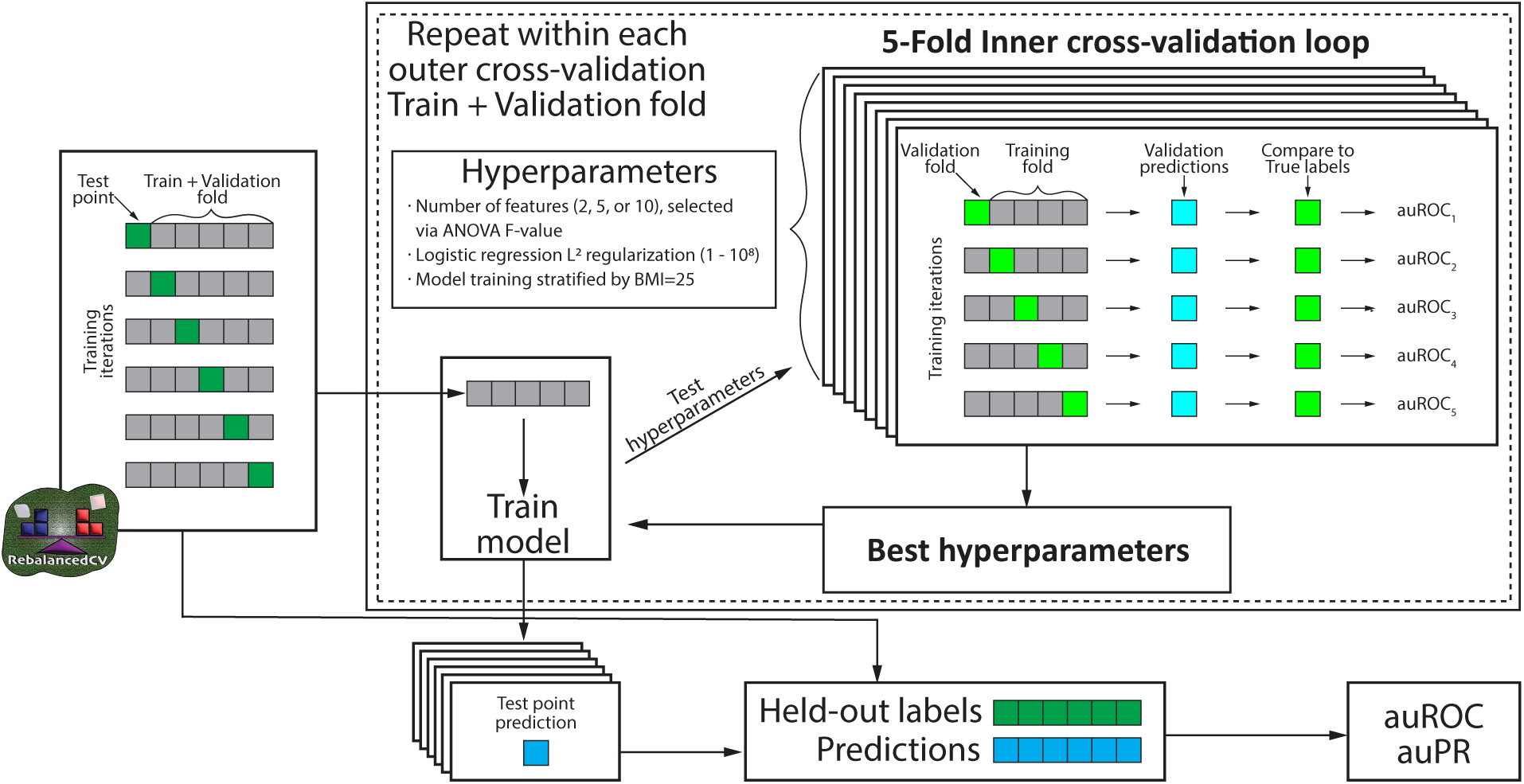
Description of our nested cross-validation pipeline. The scheme used to evaluate predictive performance within the nuMoM2b dataset, using an inner 5-fold cross-validation structure to identify optimal hyperparameters, as described in **Methods**, with the ‘Best hyperparameters’ used to train a model which implements a prediction for its corresponding outer fold left-out point.

## References

1. Duley, L. The global impact of pre-eclampsia and eclampsia. Semin. Perinatol. 33, (2009).

2. Pribadi, A. et al. Assessing the Impact of the Zero Mother Mortality Preeclampsia Program on Maternal Mortality Rates at a Single Center in Bandung, West Java (2015–2022): A Retrospective Study. Med. Sci. Monit. 29, e941097-1 (2023).

3. Hodgins, S. Pre-eclampsia as Underlying Cause for Perinatal Deaths: Time for Action. Glob Health Sci Pract 3, 525–527 (2015).

4. Chappell, L. C., Cluver, C. A., Kingdom, J. & Tong, S. Pre-eclampsia. Lancet 398, 341–354 (2021).

5. Wright, D. et al. Aspirin for Evidence-Based Preeclampsia Prevention trial: influence of compliance on beneficial effect of aspirin in prevention of preterm preeclampsia. Am. J. Obstet. Gynecol. 217, 685.e1–685.e5 (2017).

6. Phipps, E. A., Thadhani, R., Benzing, T. & Karumanchi, S. A. Pre-eclampsia: pathogenesis, novel diagnostics and therapies. Nat. Rev. Nephrol. 15, 275–289 (2019).

7. American College of Obstetricians and Gynecologists’ Committee on Practice Bulletins-Obstetrics. Gestational hypertension and preeclampsia: ACOG Practice Bulletin, number 222. Obstet. Gynecol. 135, e237–e260 (2020).

8. Zeisler, H. et al. Predictive Value of the sFlt-1:PlGF Ratio in Women with Suspected Preeclampsia. N. Engl. J. Med. 374, 13–22 (2016).

9. Rasmussen, M. et al. RNA profiles reveal signatures of future health and disease in pregnancy. Nature 601, 422–427 (2022).

10. Black, K. D. & Horowitz, J. A. Inflammatory Markers and Preeclampsia: A Systematic Review. Nurs. Res. 67, 242–251 (2018).

11. Harmon, A. C. et al. The role of inflammation in the pathology of preeclampsia. Clin. Sci. 130, 409– 419 (2016).

12. Spence, T. et al. Maternal Serum Cytokine Concentrations in Healthy Pregnancy and Preeclampsia. J. Pregnancy 2021, 6649608 (2021).

13. Prins, J. R. et al. Preeclampsia is associated with lower percentages of regulatory T cells in maternal blood. Hypertens. Pregnancy 28, 300–311 (2009).

14. Murray, E. J., Gumusoglu, S. B., Santillan, D. A. & Santillan, M. K. Manipulating CD4+ T Cell Pathways to Prevent Preeclampsia. Frontiers in Bioengineering and Biotechnology 9, (2022).

15. Aggarwal, R. et al. Association of pro- and anti-inflammatory cytokines in preeclampsia. J. Clin. Lab. Anal. 33, e22834 (2019).

16. Weel, I. C. et al. Association between Placental Lesions, Cytokines and Angiogenic Factors in Pregnant Women with Preeclampsia. PLoS One 11, e0157584 (2016).

17. Milosevic-Stevanovic, J. et al. Number of decidual natural killer cells & macrophages in pre-eclampsia. Indian J. Med. Res. 144, 823–830 (2016).

18. Kell, D. B. & Kenny, L. C. A Dormant Microbial Component in the Development of Preeclampsia. Front. Med. 3, 60 (2016).

19. Beckers, K. F. & Sones, J. L. Maternal microbiome and the hypertensive disorder of pregnancy, preeclampsia. Am. J. Physiol. Heart Circ. Physiol. 318, H1–H10 (2020).

20. Conde-Agudelo, A., Villar, J. & Lindheimer, M. Maternal infection and risk of preeclampsia: systematic review and metaanalysis. Am. J. Obstet. Gynecol. 198, 7–22 (2008).

21. Goldenberg, R. L., Hauth, J. C. & Andrews, W. W. Intrauterine infection and preterm delivery. N. Engl. J. Med. 342, 1500–1507 (2000).

22. Fettweis, J. M. et al. The vaginal microbiome and preterm birth. Nat. Med. 25, 1012–1021 (2019).

23. Berard, A. R. et al. Vaginal epithelial dysfunction is mediated by the microbiome, metabolome, and mTOR signaling. Cell Rep. 42, 112474 (2023).

24. Kindinger, L. M. et al. Relationship between vaginal microbial dysbiosis, inflammation, and pregnancy outcomes in cervical cerclage. Sci. Transl. Med. 8, 350ra102 (2016).

25. Paramel Jayaprakash, T., et al. High diversity and variability in the vaginal microbiome in women following preterm premature rupture of membranes (PPROM): A prospective cohort study. PLoS One 11, e0166794 (2016).

26. Brown, R. G. et al. Establishment of vaginal microbiota composition in early pregnancy and its association with subsequent preterm prelabor rupture of the fetal membranes. Transl. Res. 207, 30– 43 (2019).

27. Lin, C.-Y. et al. Severe preeclampsia is associated with a higher relative abundance of Prevotella bivia in the vaginal microbiota. Sci. Rep. 10, 18249 (2020).

28. Geldenhuys, J. et al. Diversity of the gut, vaginal and oral microbiome among pregnant women in South Africa with and without pre-eclampsia. Front Glob Womens Health 3, 810673 (2022).

29. Haas, D. M. et al. A description of the methods of the Nulliparous Pregnancy Outcomes Study: monitoring mothers-to-be (nuMoM2b). Am. J. Obstet. Gynecol. 212, 539.e1–539.e24 (2015).

30. Fingar, K. R. et al. Delivery hospitalizations involving preeclampsia and eclampsia, 2005–2014. in Healthcare Cost and Utilization Project (HCUP) Statistical Briefs (Agency for Healthcare Research and Quality (US), Rockville (MD), 2006).

31. Kongwattanakul, K., Saksiriwuttho, P., Chaiyarach, S. & Thepsuthammarat, K. Incidence, characteristics, maternal complications, and perinatal outcomes associated with preeclampsia with severe features and HELLP syndrome. Int. J. Womens Health 10, 371–377 (2018).

32. Yang, Y. et al. Preeclampsia Prevalence, Risk Factors, and Pregnancy Outcomes in Sweden and China. JAMA Network Open 4, e218401 (2021).

33. O’Brien, T. E., Ray, J. G. & Chan, W.-S. Maternal Body Mass Index and the Risk of Preeclampsia: A Systematic Overview. Epidemiology 14, 368 (2003).

34. Bodnar, L. M., Ness, R. B., Markovic, N. & Roberts, J. M. The risk of preeclampsia rises with increasing prepregnancy body mass index. Ann. Epidemiol. 15, 475–482 (2005).

35. Beigi, R. H., Yudin, M. H., Cosentino, L., Meyn, L. A. & Hillier, S. L. Cytokines, pregnancy, and bacterial vaginosis: comparison of levels of cervical cytokines in pregnant and nonpregnant women with bacterial vaginosis. J. Infect. Dis. 196, 1355–1360 (2007).

36. Marangoni, A. et al. New Insights into Vaginal Environment During Pregnancy. Frontiers in Molecular Biosciences 8, (2021).

37. Zanotta, N. et al. Cervico-vaginal secretion cytokine profile: A non-invasive approach to study the endometrial receptivity in IVF cycles. Am. J. Reprod. Immunol. 81, e13064 (2019).

38. Imseis, H. M. et al. Characterization of the inflammatory cytokines in the vagina during pregnancy and labor and with bacterial vaginosis. J. Soc. Gynecol. Investig. 4, 90–94 (1997).

39. Sanchez, A. et al. Variations in cytokine profiles between parous and nulliparous adult women. GREM - Gynecological and Reproductive Endocrinology & Metabolism 039–044 (2022).

40. Kumar, M. et al. Vaginal Microbiota and Cytokine Levels Predict Preterm Delivery in Asian Women. Front. Cell. Infect. Microbiol. 11, (2021).

41. Guermonprez, P. et al. Inflammatory Flt3L is essential to mobilize dendritic cells and for T cell responses during Plasmodium infection. Nat. Med. 19, 730–738 (2013-6).

42. Karsunky, H., Merad, M., Cozzio, A., Weissman, I. L. & Manz, M. G. Flt3 ligand regulates dendritic cell development from Flt3+ lymphoid and myeloid-committed progenitors to Flt3+ dendritic cells in vivo. J. Exp. Med. 198, 305–313 (2003).

43. Holgate, S. T. Epithelial damage and response. Clin. Exp. Allergy 30 Suppl 1, 37–41 (2000).

44. Cao, S. et al. Epidermal growth factor receptor activation is essential for kidney fibrosis development. Nat. Commun. 14, 7357 (2023).

45. Arend, W. P. The balance between IL-1 and IL-1Ra in disease. Cytokine Growth Factor Rev. 13, 323– 340 (2002).

46. Gabay, C., Lamacchia, C. & Palmer, G. IL-1 pathways in inflammation and human diseases. Nat. Rev. Rheumatol. 6, 232–241 (2010).

47. Armant, D. R. et al. Reduced expression of the epidermal growth factor signaling system in preeclampsia. Placenta 36, 270–278 (2015).

48. Stefańska, K. et al. Cytokine Imprint in Preeclampsia. Front. Immunol. 12, 667841 (2021).

49. Greer, I. A., Lyall, F., Perera, T., Boswell, F. & Macara, L. M. Increased concentrations of cytokines interleukin-6 and interleukin-1 receptor antagonist in plasma of women with preeclampsia: a mechanism for endothelial dysfunction? Obstet. Gynecol. 84, 937–940 (1994).

50. Huang, L. et al. A multi-kingdom collection of 33,804 reference genomes for the human vaginal microbiome. Nat Microbiol 9, 2185–2200 (2024).

51. Martino, C., et al. A novel sparse compositional technique reveals microbial perturbations. mSystems 4, (2019).

52. Allen, N. G. et al. The vaginal microbiome in women of reproductive age with healthy weight versus overweight/obesity. Obesity 30, 142–152 (2022).

53. Emanuela, F. et al. Inflammation as a link between obesity and metabolic syndrome. J. Nutr. Metab. 2012, 476380 (2012).

54. Qing, W., Shi, Y., Zhou, H. & Chen, M. Gut microbiota dysbiosis in patients with preeclampsia: A systematic review. Medicine in Microecology 10, 100047 (2021).

55. Miao, T. et al. Decrease in abundance of bacteria of the genus Bifidobacterium in gut microbiota may be related to pre-eclampsia progression in women from East China. Food Nutr. Res. 65, (2021).

56. Schwebke, J. R., Muzny, C. A. & Josey, W. E. Role of Gardnerella vaginalis in the pathogenesis of bacterial vaginosis: a conceptual model. J. Infect. Dis. 210, 338–343 (2014).

57. Riedel, C.-U. et al. Anti-inflammatory effects of bifidobacteria by inhibition of LPS-induced NF-kappaB activation. World J. Gastroenterol. 12, 3729–3735 (2006).

58. Ruiz, L., Delgado, S., Ruas-Madiedo, P., Sánchez, B. & Margolles, A. Bifidobacteria and Their Molecular Communication with the Immune System. Front. Microbiol. 8, 2345 (2017).

59. Cornejo, O. E., Hickey, R. J., Suzuki, H. & Forney, L. J. Focusing the diversity of Gardnerella vaginalis through the lens of ecotypes. Evol. Appl. 11, 312–324 (2018).

60. Park, S. et al. Ureaplasma and Prevotella colonization with Lactobacillus abundance during pregnancy facilitates term birth. Sci. Rep. 12, 10148 (2022).

61. Pelayo, P. et al. Prevotella are major contributors of sialidases in the human vaginal microbiome. bioRxiv 2024.01.09.574895 (2024) doi:10.1101/2024.01.09.574895.

62. Segui-Perez, C. et al. Prevotella timonensis degrades the vaginal epithelial glycocalyx through high fucosidase and sialidase activities. bioRxiv 2024.01.09.574844 (2024) doi:10.1101/2024.01.09.574844.

63. Farr Zuend, C., et al. Increased genital mucosal cytokines in Canadian women associate with higher antigen-presenting cells, inflammatory metabolites, epithelial barrier disruption, and the depletion of L. crispatus. Microbiome 11, 159 (2023).

64. Anahtar, M. N. et al. Cervicovaginal bacteria are a major modulator of host inflammatory responses in the female genital tract. Immunity 42, 965–976 (2015).

65. Escalda, C., Botelho, J., Mendes, J. J. & Machado, V. Association of bacterial vaginosis with periodontitis in a cross-sectional American nationwide survey. Sci. Rep. 11, 630 (2021).

66. Keller, M. J. et al. Longitudinal assessment of systemic and genital tract inflammatory markers and endogenous genital tract E. coli inhibitory activity in HIV-infected and uninfected women. Am. J. Reprod. Immunol. 75, 631–642 (2016).

67. Singh, A. et al. DIABLO: an integrative approach for identifying key molecular drivers from multi-omics assays. Bioinformatics 35, 3055–3062 (2019).

68. Hassan, G. S., Merhi, Y. & Mourad, W. CD40 ligand: a neo-inflammatory molecule in vascular diseases. Immunobiology 217, 521–532 (2012).

69. Wood, G. W., Hausmann, E. & Choudhuri, R. Relative role of CSF-1, MCP-1/JE, and RANTES in macrophage recruitment during successful pregnancy. Mol. Reprod. Dev. 46, 62–9; discussion 69-70 (1997).

70. Mellembakken, J. R., Solum, N. O., Ueland, T., Videm, V. & Aukrust, P. Increased concentrations of soluble CD40 ligand, RANTES and GRO-alpha in preeclampsia--possible role of platelet activation. Thromb. Haemost. 86, 1272–1276 (2001).

71. Kauma, S. et al. Increased endothelial monocyte chemoattractant protein-1 and interleukin-8 in preeclampsia. Obstet. Gynecol. 100, 706–714 (2002).

72. Redman, C. W., Sacks, G. P. & Sargent, I. L. Preeclampsia: an excessive maternal inflammatory response to pregnancy. Am. J. Obstet. Gynecol. 180, 499–506 (1999).

73. Li, J. et al. Gut microbiota dysbiosis contributes to the development of hypertension. Microbiome 5, 14 (2017).

74. Dinakis, E. et al. Association between the gut microbiome and their metabolites with human blood pressure variability. Hypertension 79, 1690–1701 (2022).

75. Mikolajczyk, T. P. et al. Role of chemokine RANTES in the regulation of perivascular inflammation, T-cell accumulation, and vascular dysfunction in hypertension. FASEB J. 30, 1987–1999 (2016).

76. Rabkin, S. W., Langer, A., Ur, E., Calciu, C.-D. & Leiter, L. A. Inflammatory biomarkers CRP, MCP-1, serum amyloid alpha and interleukin-18 in patients with HTN and dyslipidemia: impact of diabetes mellitus on metabolic syndrome and the effect of statin therapy. Hypertens. Res. 36, 550– 558 (2013).

77. Tan, J. et al. Association between maternal weight indicators and iron deficiency anemia during pregnancy: A cohort study: A cohort study. Chin. Med. J. (Engl.) 131, 2566–2574 (2018).

78. Callahan, B. J. et al. Replication and refinement of a vaginal microbial signature of preterm birth in two racially distinct cohorts of US women. Proceedings of the National Academy of Sciences 114, 9966– 9971 (2017).

79. Starc, M., Lučovnik, M., Eržen Vrlič, P. & Jeverica, S. Protective effect of Lactobacillus crispatus against vaginal colonization with group B streptococci in the third trimester of pregnancy. Pathogens 11, 980 (2022).

80. Nunn, K. L. et al. Enhanced trapping of HIV-1 by human cervicovaginal mucus is associated with Lactobacillus crispatus-dominant Microbiota. MBio 6, e01084–15 (2015).

81. McCoy, Z. T. et al. Antibody Response to the Sneathia vaginalis Cytopathogenic Toxin A during Pregnancy. ImmunoHorizons 8, 114–121 (2024).

82. Theis, K. R. et al. Sneathia: an emerging pathogen in female reproductive disease and adverse perinatal outcomes. Crit. Rev. Microbiol. 47, 517–542 (2021).

83. Austin, G. I., Pe’er, I. & Korem, T. Distributional bias compromises leave-one-out cross-validation. arXiv [stat.ME] (2024).

84. Tiruneh, S. A. et al. Externally validated prediction models for pre-eclampsia: systematic review and meta-analysis. Ultrasound Obstet. Gynecol. 63, 592–604 (2024).

85. Li, S., et al. Improving preeclampsia risk prediction by modeling pregnancy trajectories from routinely collected electronic medical record data. NPJ Digital Medicine 5, (2022).

86. Lin, Y. C. et al. Preeclampsia Predictor with Machine Learning: A Comprehensive and Bias-Free Machine Learning Pipeline. medRxiv 2022.06.08.22276107 (2022) doi:10.1101/2022.06.08.22276107.

87. Ghaemi, M. S. et al. Proteomic signatures predict preeclampsia in individual cohorts but not across cohorts - implications for clinical biomarker studies. J. Matern. Fetal. Neonatal Med. 35, 5621–5628 (2022).

88. Romero, R. et al. The vaginal Microbiota of pregnant women varies with gestational age, maternal age, and parity. Microbiol. Spectr. 11, e0342922 (2023).

89. Austin, G. I. et al. Processing-bias correction with DEBIAS-M improves cross-study generalization of microbiome-based prediction models. bioRxiv 2024.02.09.579716 (2024) doi:10.1101/2024.02.09.579716.

90. Moufarrej, M. N. et al. Early prediction of preeclampsia in pregnancy with cell-free RNA. Nature 602, 689–694 (2022).

91. Lee, S. K., Kim, C. J., Kim, D.-J. & Kang, J.-H. Immune cells in the female reproductive tract. Immune Netw. 15, 16–26 (2015).

92. Smith, S. D., Dunk, C. E., Aplin, J. D., Harris, L. K. & Jones, R. L. Evidence for immune cell involvement in decidual spiral arteriole remodeling in early human pregnancy. Am. J. Pathol. 174, 1959–1971 (2009).

93. Broekhuizen, M. et al. The Placental Innate Immune System Is Altered in Early-Onset Preeclampsia, but Not in Late-Onset Preeclampsia. Front. Immunol. 12, (2021).

94. Menkhorst, E. et al. IL11 activates the placental inflammasome to drive preeclampsia. Front. Immunol. 14, 1175926 (2023).

95. Karge, A. et al. Performance of sFlt-1/PIGF ratio for the prediction of perinatal outcome in obese pre-eclamptic women. J. Clin. Med. 11, 3023 (2022).

96. Klein, K. O. et al. Effect of obesity on estradiol level, and its relationship to leptin, bone maturation, and bone mineral density in children. J. Clin. Endocrinol. Metab. 83, 3469–3475 (1998).

97. Shen, Z. et al. Effect of BMI on the value of serum progesterone to predict clinical pregnancy outcome in IVF/ICSI cycles: a retrospective cohort study. Front. Endocrinol. (Lausanne) 14, 1162302 (2023).

98. Ingram, K., Ngalame Eko, E., Nunziato, J., Ahrens, M. & Howell, B. Impact of obesity on the perinatal vaginal environment and bacterial microbiome: effects on birth outcomes. J. Med. Microbiol. 73, 001874 (2024).

99. McGregor, J. A. et al. Bacterial vaginosis is associated with prematurity and vaginal fluid mucinase and sialidase: results of a controlled trial of topical clindamycin cream. Am. J. Obstet. Gynecol. 170, 1048–59; discussion 1059-60 (1994).

100. Cauci, S. & Culhane, J. F. High sialidase levels increase preterm birth risk among women who are bacterial vaginosis-positive in early gestation. Am. J. Obstet. Gynecol. 204, 142.e1–9 (2011).

101. Lin, C.-Y. et al. Severe preeclampsia is associated with a higher relative abundance of Prevotella bivia in the vaginal microbiota. Sci. Rep. 10, 18249 (2020).

102. Freitas, A. C. & Hill, J. E. Quantification, isolation and characterization of Bifidobacterium from the vaginal microbiomes of reproductive aged women. Anaerobe 47, 145–156 (2017).

103. Schwecht, I., Nazli, A., Gill, B. & Kaushic, C. Lactic acid enhances vaginal epithelial barrier integrity and ameliorates inflammatory effects of dysbiotic short chain fatty acids and HIV-1. Sci. Rep. 13, 20065 (2023).

104. Romero, R. et al. Sterile and microbial-associated intra-amniotic inflammation in preterm prelabor rupture of membranes. J. Matern. Fetal. Neonatal Med. 28, 1394–1409 (2015).

105. Srinivasan, S. et al. Bacterial communities in women with bacterial vaginosis: high resolution phylogenetic analyses reveal relationships of microbiota to clinical criteria. PLoS One 7, e37818 (2012).

106. Ali, A. A., Rayis, D. A., Abdallah, T. M., Elbashir, M. I. & Adam, I. Severe anaemia is associated with a higher risk for preeclampsia and poor perinatal outcomes in Kassala hospital, eastern Sudan. BMC Res. Notes 4, 311 (2011).

107. Johnson, A., Vaithilingan, S. & Avudaiappan, S. L. The interplay of hypertension and anemia on pregnancy outcomes. Cureus 15, e46390 (2023).

108. Rabkin, S. W. The role of interleukin 18 in the pathogenesis of hypertension-induced vascular disease. Nat. Clin. Pract. Cardiovasc. Med. 6, 192–199 (2009).

109. Ferroni, P. & Guadagni, F. Soluble CD40L and its role in essential hypertension: diagnostic and therapeutic implications. Cardiovasc. Hematol. Disord. Drug Targets 8, 194–202 (2008).

110. Lu, X. et al. Classical Dendritic Cells Mediate Hypertension by Promoting Renal Oxidative Stress and Fluid Retention. Hypertension 75, 131–138 (2020).

111. Stefan, N., Häring, H.-U., Hu, F. B. & Schulze, M. B. Metabolically healthy obesity: epidemiology, mechanisms, and clinical implications. Lancet Diabetes Endocrinol. 1, 152–162 (2013).

112. Greenland, P. et al. Large-Scale Proteomics in Early Pregnancy and Hypertensive Disorders of Pregnancy. JAMA Cardiol (2024) doi:10.1001/jamacardio.2024.1621.

113. Hypertension in pregnancy. Report of the American College of Obstetricians and Gynecologists’ Task Force on Hypertension in Pregnancy. Obstet. Gynecol. 122, 1122–1131 (2013).

114. ACOG Committee Opinion No. 743 Summary: Low-Dose Aspirin Use During Pregnancy. Obstet. Gynecol. 132, 254–256 (2018).

115. Bolger, A. M., Lohse, M. & Usadel, B. Trimmomatic: a flexible trimmer for Illumina sequence data. Bioinformatics 30, 2114–2120 (2014).

116. Langmead, B. & Salzberg, S. L. Fast gapped-read alignment with Bowtie 2. Nat. Methods 9, 357–359 (2012).

117. Li, H. et al. The Sequence Alignment/Map format and SAMtools. Bioinformatics 25, 2078–2079 (2009).

118. Wood, D. E., Lu, J. & Langmead, B. Improved metagenomic analysis with Kraken 2. Genome Biol. 20, 257 (2019).

119. Lu, J., Breitwieser, F. P., Thielen, P. & Salzberg, S. L. Bracken: estimating species abundance in metagenomics data. PeerJ Comput. Sci. 3, e104 (2017).

120. Pedregosa, F. et al. Scikit-learn: Machine Learning in Python. J. Mach. Learn. Res. 12, 2825–2830 (2011).

121. Mason, S. J. & Graham, N. E. Areas beneath the relative operating characteristics (ROC) and relative operating levels (ROL) curves: Statistical significance and interpretation. Q. J. R. Meteorol. Soc. 128, 2145–2166 (2002).

